# Ndr/Lats kinases bind specific Mob-family coactivators through a conserved and modular interface

**DOI:** 10.1101/242115

**Authors:** Benjamin W. Parker, Gergő Gógl, Mónika Bálint, Csaba Hetenyi, Attila Remenyi, Eric L. Weiss

## Abstract

Ndr/Lats kinases bind Mob coactivator proteins to form complexes that are essential and deeply conserved components of “Hippo” signaling pathways, which control cell proliferation and morphogenesis in eukaryotes. All Ndr/Lats kinases have a characteristic N-terminal region (NTR) that binds a specific Mob co-factor: Lats kinases associate with Mob1 proteins, and Ndr kinases associate with Mob2 proteins. To better understand the functional significance of Mob protein association with Ndr/Lats kinases and selective binding of Ndr and Lats to distinct Mob co-factors, we solved crystal structures of *Saccharomyces cerevisiae* Cbk1(NTR)-Mob2 and Dbf2(NTR)-Mob1 and experimentally assessed determinants of Mob cofactor binding and specificity. This significantly refines the previously determined structure of Cbk1 kinase bound to Mob2, presently the only crystallographic model of a full length Ndr/Lats kinase complexed with a Mob cofactor. Our analysis indicates that the Ndr/Lats NTR-Mob interface provides a distinctive kinase regulation mechanism, in which Mob co-factor organizes the Ndr/Lats NTR to interact with the AGC kinase C-terminal hydrophobic motif (HM) activation segment. The Mob-organized NTR appears to mediate HM association with an allosteric site on the kinase N-lobe. We also found that Cbk1 and Dbf2 associated highly specifically with Mob2 and Mob1, respectively. Alteration of specific positions in the Cbk1 NTR allows association of non-cognate Mob co-factor, indicating that cofactor specificity is restricted by discrete sites rather than broadly distributed. Overall, our analysis provides a new picture of the functional role of Mob association and indicates that the Ndr/Lats(NTR)-Mob interface overall is largely a common structural platform that mediates kinase-cofactor binding.

## Introduction

“Hippo” signaling systems are widespread in the eukaryotic world, playing roles in cell proliferation control, tissue development, and cell morphogenesis (1). These pathways have a deeply conserved signaling core in which upstream Ste20-family Mst/*hippo* kinases phosphorylate key C-terminal activating sites of Ndr/Lats kinases, which belong to the AGC superfamily of protein kinases. The Ndr/Lats kinases fall into Ndr and Lats subfamilies that, while closely related, are distinct from one another in amino acid sequence and cellular function from yeast to animals (reviewed in (2–4)). In animals, for example, Lats-related kinases inhibit YAP/TAZ family transcriptional coactivators driving genes that promote cell cycle entry and apoptosis resistance; this system links cell proliferation to tissue organization (1,4–6). Animal Ndr kinases play separate and incompletely understood roles in cell proliferation and morphogenesis of polarized structures, notably organization of neurites (1–5).

Ndr/Lats kinases have a distinctive segment immediately N-terminal to their kinase domains generally referred to as the NTR (N-terminal regulatory) region. This NTR region binds to a small “Mob” coactivator protein, an association that appears to be essential for kinase function (4,7,8). In the budding yeast *S. cerevisiae*, where Mob coactivators were discovered, the paralogous Lats-related kinases Dbf2 and Dbf20 bind the Mob1 protein, forming a central component of a hippo pathway system called the mitotic exit network (MEN) that controls cytokinesis and the transition from M phase to G1 (9–20) (reviewed in (21,22)). The budding yeast Ndr-subfamily kinase Cbk1 associates with the Mob2 coactivator and functions in a different pathway that controls the final stage of cell separation and polarized cell growth called the Regulation of Ace2 and Morphogenesis (RAM) network (23–26).

Notably, the MEN and the RAM network control distinct processes that are important for late cell division and morphogenesis, with no evidence that Dbf2/20 and Cbk1 have overlapping functions (compare time courses shown in (10–12,26)). For example, the phosphatase Cdc14 is functionally downstream of Dbf2-Mob1 and the MEN (27), but functionally upstream of Cbk1-Mob2 and the RAM network (26,28). Additionally, the RAM network promotes sustained polarized growth and controls the activity of a translation-regulating mRNA binding protein (26,29), while the MEN promotes downregulation of Cdk-targets and cytokinesis (22,30). Notably, Cbk1-Mob1 or Dbf2/20-Mob2 complexes do not appear to form (23,24,26,31–38), despite simultaneous presence of all of the proteins in the cytosol, suggesting a mechanism that enforces kinase-coactivator association specificity. Studies of *Schizosaccharomyces pombe* Sid2 (Lats)-Mob1 and Orb6 (Ndr)-Mob2 further indicate highly specific Ndr/Lats-Mob interactions (39), despite generally high conservation of Mob cofactors and the kinase NTR region across the entire Ndr/Lats family (40).

In animals, biochemical, genetic, and *in vivo* analyses indicate that Lats kinases associate with Mob1 proteins and Ndr kinases associate with Mob2 proteins as functional units (40). However, human Ndr kinases may also associate with both Mob1 and Mob2. Under certain conditions Ndr1/2 and Mob1 co-immunoprecipitate, and the proteins been linked to one another in proteome-scale affinity purification mass spec (AP-MS) experiments (41,42). Moreover, Kulaberoglu et al. recently provided a crystal structure of an *in vitro* assembled Ndr2(NTR fragment)-Mob1A complex (43). It remains unclear if such Ndr-Mob1 complexes are physiologically significant, but characterization of Ndr kinase functions in human cells is currently in early stages.

Mob coactivator binding is necessary for Ndr/Lats kinase function, but the mechanism underlying this Mob-mediated kinase activation is not well understood. A structure of the Cbk1-Mob2 complex contributed by our groups, currently the only crystallographic model of a full-length Ndr/Lats kinase bound to a Mob coactivator, demonstrates that the Mob-bound NTR region and the kinase N-lobe form a dramatic cleft. This cleft associates with Cbk1’s C-terminal hydrophobic motif (HM) region, a phosphorylation-regulated activation module that is broadly present in the large AGC kinase family. Unfortunately, low resolution in this structure at the kinase-Mob interface significantly complicates analysis of the function of Mob binding. Additional crystallographic studies of NTR region fragments of human Lats1 and Ndr1 bound to Mob1 have provided substantially better assessment of Mob coactivator binding (43–45). Key questions remain, however, about structural organization of the full-length kinase-coactivator complex, the Mob protein’s role in kinase activation, and Mob cofactor binding specificity.

Here we describe new crystallographic analysis of a Cbk1(NTR fragment)-Mob2 complex that provides a higher resolution view of the Cbk1-Mob2 interface. This structure allowed us to reanalyze the lower resolution electron density map of the full Cbk1-Mob2 complex (46), showing that Cbk1’s NTR adopts a dramatic two-helix hairpin in the full kinase-coactivator complex that is highly similar to structures of human Lats1(NTR fragment)-Mob1 (44,45) and Ndr1(NTR fragment)-Mob1 (43). The new structure also provides more precise positioning of Cbk1’s C-terminal HM region relative to the kinase’s N-lobe and NTR region. In this view, Mob association with the NTR region produces a structural system that coordinates the key activating HM phosphorylation site with a highly conserved positively charged amino acid in the NTR region. Electrostatic association of these positions would allow HM region aromatic side chains to promote ordering of the kinase’s otherwise flexible C helix, which is critical for kinase activation. Our analyses further shows that Mob1 binds Dbf2^NTR^ and Mob2 binds Cbk1^NTR^ with both high specificity and affinity. In addition to highlighting conserved mechanisms by which Mob proteins bind Ndr/Lats kinases, our analysis identified a Mob protein tripeptide at the Mob-NTR region interface that differs between Mob1 and Mob2 family proteins and contributes strongly to specificity of kinase – cofactor association. This short region functions as a “Kinase Restrictor Motif” in yeast Mob proteins by greatly reducing their association with off target Ndr/Lats kinase NTR regions.

## Results

### A Cbk1 (NTR fragment) - Mob2 structure significantly improves analysis of multiple kinase-coactivator features

As noted, mechanisms by which Mob cofactors bind Ndr/Lats family kinases and regulate their activation was complicated by uncertain electron density in the kinase-Mob coactivator association region of Cbk1-Mob2, the sole structure of a full length Ndr/Lats kinase bound to a Mob coactivator. We therefore sought to improve the resolution of this interface through co-crystallization of the N-terminal region (NTR) of Cbk1 (Cbk1^NTR^) with Mob2. Consistent with prior report (17), we were unable to stably express monomeric Mob2 in *E.coli*. We therefore engineered a Mob2 ^V148C Y153C^ allele that replaces a zinc-binding motif lost in budding yeast Mob2 but found in most metazoan Mob2 orthologs including *S. pombe* Mob2 and *S. cerevisiae* Mob1. This change stabilized Mob2 and allowed suitable bacterial expression for biochemistry; notably, the reintroduced zinc binding site is distant from the Mob2 surface that interacts with Cbk1’s NTR region. Except where specifically noted, Mob2 constructs described here carry the Mob2 ^V148C Y153C^ substitutions. We crystallized Mob2^45-287^ bound to Cbk1^NTR^ (Cbk1^251-351^) to 2.8 Å resolution (Table 1 and Figure 1A). This higher resolution Cbk1^NTR^-Mob2 data allowed us to revisit prior crystallographic analysis and modeling of the full-length Cbk1-Mob2 complex that resulted in significant improvement of the Cbk1-Mob2 complex electron density map.

**Figure 1.**
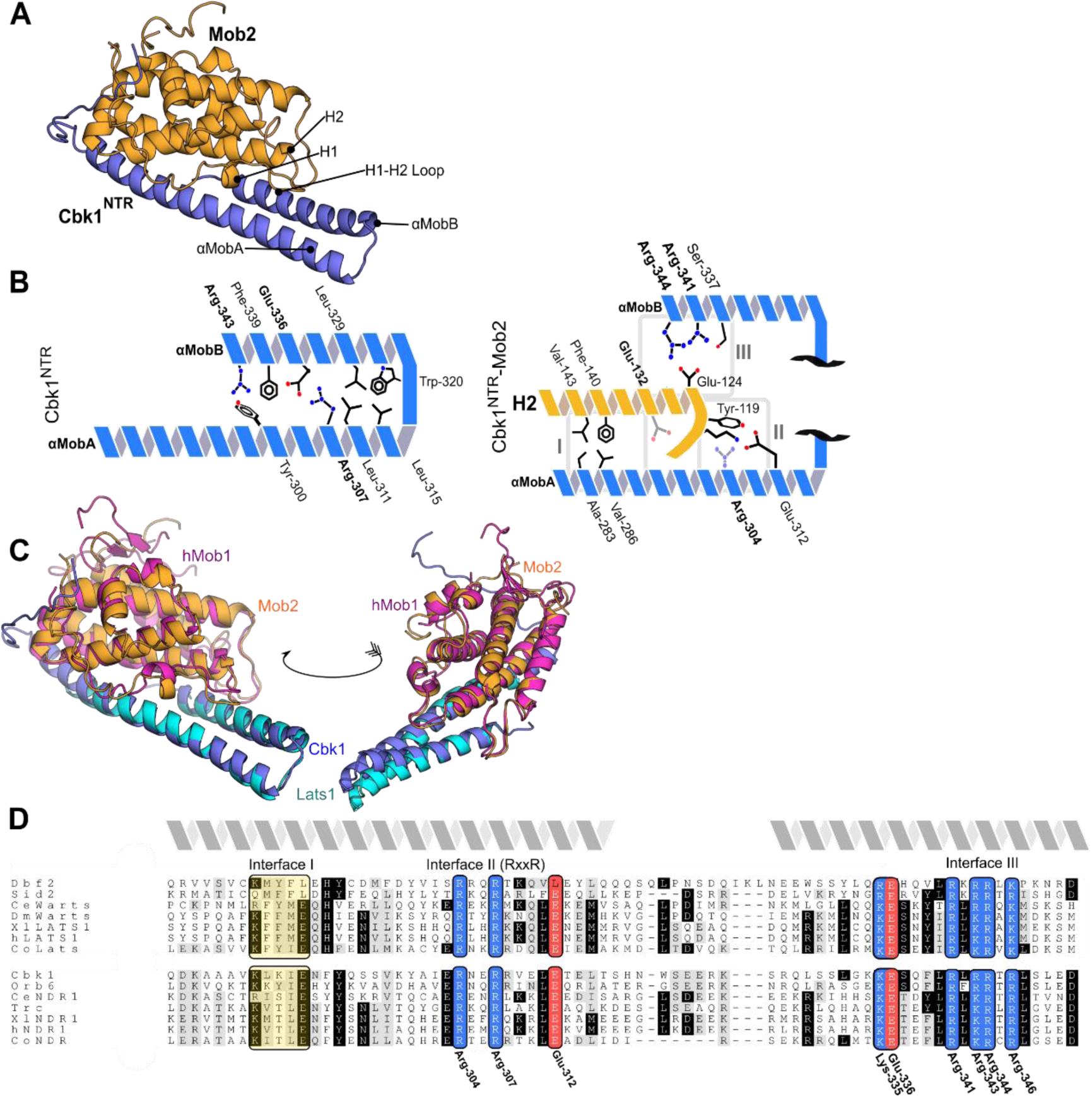
The Cbk1 ^NTR^ -Mob2 complex contains conserved structural motifs. (A) Cbk1 ^NTR^ (251-351, blue) complexed with Mob2 ^V148C Y153C^ (45-287, orange). Mob-family helices and motifs are highlighted. (B) The revised Cbk1-Mob2 interaction architecture. Conserved residues are bolded. The left panel illustrates amino acids likely mediating the αMobA and B interaction. The right panel illustrates interactions with the Mob2 H2 helix. Canonical interfaces I-III between the Mob and the Ndr/Lats kinase are boxed. (C) Cbk1 ^NTR^ -Mob2 (blue/orange) overlaid with Lats1 ^NTR^ - hMob1 (PDB code 5B5W, cyan/pink). (D) Alignment of the Ndr/Lats ^NTR^ regions across eukaryotes. Ce, *Caenorhabditis elegans*; Dm, *Drosophila melanogaster*; Xl, *Xenopus laevis*; Co, *Capsaspora owczarzaki*. Basic residues, blue; acidic residues, red; hydrophobic regions, yellow. Interfaces I-III equivalent to those described in the Lats1-hMob1 structure are labeled. Residues conserved with Cbk1 are labeled.

**Table 1.** Crystallographic data collection and refinement

|  | Dbf2<sup>NTR</sup>- Mob1 | Cbk1<sup>NTR</sup>- Mob2 | Cbk1- Mob2 -<br>pepSsd1 |
| --- | --- | --- | --- |
| Data collection |  |  |  |
| Wavelength (Å) | 0.978560 | 0.978720 | 1.0000 |
| Space group | P 6<sub>1</sub> 2 2 | P 4<sub>1</sub> 2<sub>1</sub> 2 | C 1 2 1 |
| Cell dimensions |  |  |  |
| *a*, *b*, *c* (Å) | 108.28, 108.28, 134.76 | 126.23, 126.27, 49.34 | 138.43, 79.99, 117.59 |
| $\alpha$, $\beta$, $\gamma$ (°) | 90.0, 90.0, 120.0 | 90.0, 90.0, 90.0 | 90.0, 117.6, 90.0 |
| Resolution (Å) | 46.9-3.5 (3.59-3.50) | 39.9-2.8 (2.87-2.80) | 44.60-3.15 (3.23-3.15) |
| CC<sub>1/2</sub> | 99.0 (79.1) | 100.0 (67.0) | 99.6 (80.4) |
| *R*<sub>merge</sub> | 54.1 (259.2) | 10.0 (152.4) | 16.8 (130.4) |
| Mean *I*/σ*I* | 9.34 (1.49) | 21.79 (2.21) | 9.37 (2.05) |
| Completeness (%) | 99.9 (100) | 99.8 (100) | 99.7 (99.7) |
| Redundancy | 24.60 (25.83) | 14.16 (14.74) | 7.73 (7.82) |
| No. reflections | 6385 (447) | 10323 (750) | 19888 (1455) |
| Refinement |  |  |  |
| *R*<sub>work</sub>/*R*<sub>free</sub> | 0.2292/0.2631 | 0.2490/0.2838 | 0.2310/0.2983 |
| No. atoms |  |  |  |
| Protein | 2024 | 2196 | 4715 |
| Ligand/ion | 17 | 1 | 31 |
| B-factors |  |  |  |
| Protein | 88.3 | 92.7 | 98.8 |
| Ligand | 139.1 | 114.7 | 92.6 |
| Ramachandran |  |  |  |
| Favored (%) | 93.5 | 93 | 76** |
| Allowed (%) | 5 | 4 | 23** |
| Outliers (%) | 1.6 | 3 | 1** |
| Rotamer outliers (%) | 1.85 | 15 | 21.5 |
| R.m.s deviations |  |  |  |
| Bond lengths (Å) | 0.010 | 0.004 | 0.011 |
| Bond angles (°) | 0.75 | 0.70 | 1.35 |

\*Highest resolution shell is shown in parenthesis.
\*\*Ramachandran restraints were used during refinement.

In the Cbk1^NTR^-Mob2 complex the kinase NTR region forms a dramatic V-shaped arrangement of two helices (Figure 1A). Importantly, this NTR region structure is also strongly supported in the improved electron density map of the full length Cbk1-Mob2 complex; we have revised incorrect threading of this region in PDB (PDB 5NCL) (Supplementary Figure 1). The updated Cbk1-Mob2 structure highlights residues responsible for cohesion of the NTR region αMob helices to each other (Figure 1B, left) and adhesion of the NTR region to the Mob2 cofactor, specifically helix H2 of Mob2 (Figure 1B, right). Importantly, the bi-helical hairpin organization of Cbk1’s NTR region in Mob2 complexes is largely superimposable with recent structures of metazoan Ndr/Lats NTR fragments complexed with Mob1 (43–45). Overlay of Cbk1^NTR^ -Mob2 with the Lats1 ^NTR^ -hMob1 (PDB: 5B5W) crystal structure shows striking similarity (Figure 1C; RMSD 1.082Å). Regions identified as important for binding are highly conserved across eukaryotes (Figure 1D), with interfaces II and III showing similarity in the alignment. This includes conserved Cbk1 residues Arg-307 (Interface II) which has a similar geometry to Lats1 Arg-660 (44,45) and mediates the interaction with αMobB Glu-336. Furthermore, Cbk1 Arg-344 and Arg-341 interact with Mob2 Glu-124 in interface III. Overall, this analysis provides a picture of a kinase-coactivator interface that is structurally homologous across vast evolutionary distances in eukaryotes.

Improvement of the full length Mob2-Cbk1 electron density map allowed significant revision of structural features with that were previously ambiguous as well as description of additional ones. These include conformational organization of the kinase’s DFG motif, which is an important part of the enzyme’s activation mechanism, as well as resolution of an N-terminal extension of Cbk1 (Supplementary Figure 4). Our previously reported structure of full length Cbk1-Mob2 used kinase-coactivator crystals grown in the presence of a short kinase domain docking peptide from Ssd1, a conserved RNA binding protein that represses translation of specific mRNAs (29,47). This docking peptide is functionally important in Cbk1’s negative regulation of Ssd1 (46). Electron density corresponding to the Ssd1 peptide is not convincingly evident in the previously reported crystal structure. Our improved map, however, clearly shows additional electron density close to a putative docking peptide binding groove previously supported by both computational and mutational analysis (Figure 2A, inset, bottom). This observed density is close to Phe-447, Trp-444 and Tyr-687, which are important for Ssd1 peptide binding, but not close to Phe-699, which was not required for peptide binding (46). These data allow direct crystallographic mapping of docking peptide association with the Cbk1 kinase domain, corroborating and moderately refining the peptide conformation suggested by combined mutational and computational approaches.

**Figure 2.**
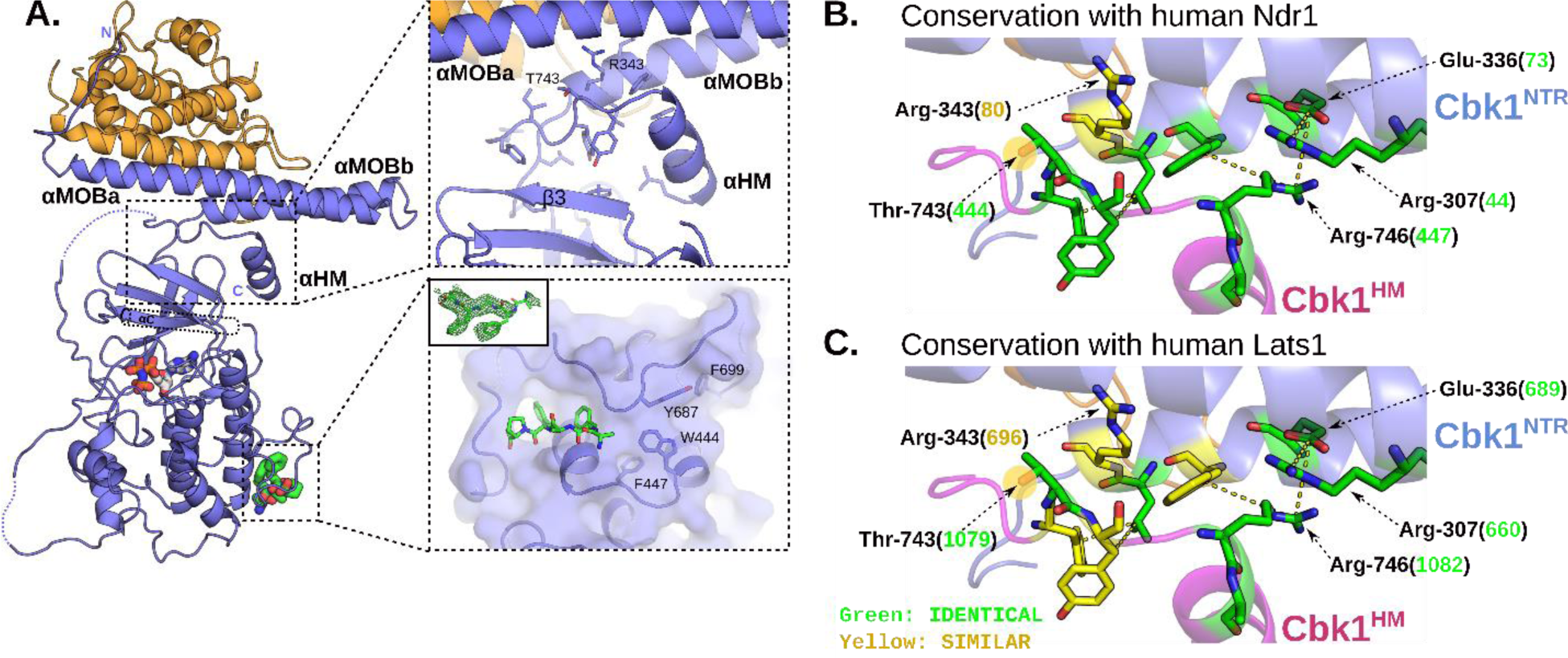
The revised Cbk1-Mob2 crystal structure reveals novel Ndr structural components conserved in Ndr1/Lats1. (A) The crystal structure of the inactive Cbk1-Mob2 complex was remodeled based on the new Cbk1 ^NTR^ -Mob2 structure. In the new model the coactivator binds to two N-terminal helices (αMobA and αMobB) which together with the top of the kinase domain (β3) form a binding slot for the AGC kinase hydrophobic motif (HM). The C-terminal part of this motif also adopts a helical conformation (αHM). The phosphorylation site on the HM (Thr-743) is next to Arg-343 from αMobB, hinting on the importance of Mob coactivator binding in the allosteric phosphoregulation of the Ndr/Lats kinase domain. Crystallographic model rebuilding also improved the overall quality of the electron density map which enabled us to trace the Ssd1 docking peptide (green sticks). The inset shows the simulated annealing Fo-Fc omit map contoured at 2 σ. (B) and (C) Residues in the interacting region between the HM and the Cbk1^NTR^ are conserved in human Ndr1/Lats1. Green residues are identical; yellow are chemically similar. Numbers in parenthesis indicate the human Ndr/Lats residue orthologous to the indicated Cbk1 amino acid (black). (B) Major interacting residues between Cbk1^NTR^ and the Cbk1 hydrophobic motif are compared with orthologues in human Ndr1. (C) The NTR-HM interface is compared with human Lats1.

Most notably, enhancement of the Cbk1-Mob2 structure resolved the C-terminal extension of Cbk1’s hydrophobic motif (HM), providing a new view of the conformation of this important activating segment. Cbk1’s C-terminal HM regulatory region is helical (denoted αHM) in the revised structure, and contacts residues in Cbk1’s αMobB and the core of the kinase domain (β3) (Figure 2A, inset, top). We also found that Arg-746 in Cbk1’s αHM interacts with Cbk1 Glu-336, which is positioned between the kinase NTR region’s αMob helices. This is consistent with notable phosphorylation-independent and non-activating interaction of the HM with the kinase domain we previously described (Figures 2B and C) (46). Phosphorylation of Thr-743 in Cbk1’s C-terminal HM is a crucial activating modification; a similar regulatory site is broadly present in AGC kinases. Notably, positioning of Cbk1’s HM relative to a highly conserved arginine in the NTR region (Arg-343, Figure 2A, inset, top see also Figure 2B and C) suggests that activating phosphorylation of Thr-743 provides an electrostatic interaction with Arg-343 that stabilizes the conformation of the HM, allowing an engagement of its aromatic side chains with the kinase N-lobe that drives disorder-to-order transition of the crucial kinase subdomain C helix (Figure 3). This suggests an activating phosphorylation at Thr-743 could hypothetically provide an electrostatic interaction to stabilize the HM conformation, which in turn may drive a disorder-to-order transition of the crucial kinase subdomain C helix similar to that observed in the related PKB mechanism of activation (48). This activation model suggests that coactivator binding is not only essential in HM binding but also in the precise coordination of the kinase’s active state when it is phosphorylated.

**Figure 3.**
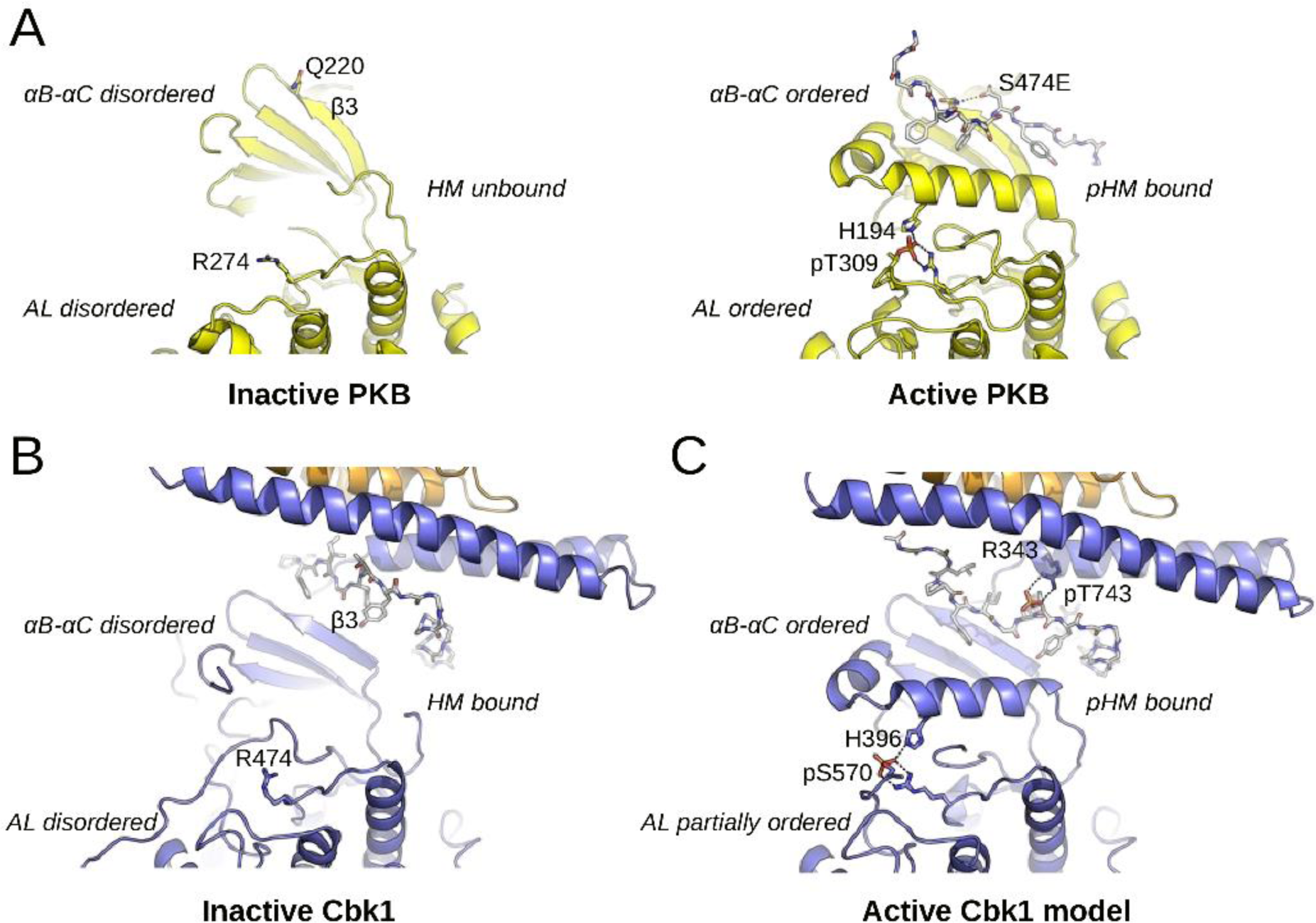
Cbk1 activation: the role of HM phosphorylation and Mob binding. (A) The activation of PKB involves a disorder-to-order transition at the αB-αC segment, the activation loop (AL) and the HM. (B) The inactive Cbk1 structure shows a similar structure apart from the position of its HM. The inactive crystal structure of the Cbk1-Mob2 shows that the HM is bound to the groove formed by the Mob bound NTR and β3 from Cbk1. (C) For activation, the ordering of αC helix in Cbk1 similarly to PKB is likely required (MD model). This latter may be supported by interactions formed by phospho-HM and phospho-AL. The activation loop phosphosite (PKB: Thr-309, Cbk1: Ser-570) is coordinated by an arginine residue from the kinase HRD motif (PKB: Arg-274, Cbk1: Arg-474) and by another residue from the αC helix (PKB: His-194, Cbk1: His-396). The hydrophobic motif phosphosite (PKB: Ser-474, Cbk1: Thr-743) in most AGC kinases is coordinated by a single residue from β3 (PKB: Gln-220). In the case of Cbk1 an arginine residue (Arg-343) from the Mob bound αMobB coordinates the HM phosphosite while β3 may not be involved. Both for PKB and Cbk1 phospho-HM coordination (by Gln-220-pSer-474 or by Arg-343-pThr-743) bring hydrophobic residues much closer to the αC and probably facilitates its ordering. This activation model suggests that coactivator binding is not only essential in HM binding but also in the precise coordination of the kinase’s active state when it is phosphorylated. PKB is colored in yellow and both HM regions are shown with gray sticks.

Strikingly, all major interacting amino acids we identified in Cbk1’s NTR and HM are either identical or highly similar at corresponding sites in human Lats1 and Ndr1 (Figure 2B and C). The crucial Arg-343 of Cbk1 is charge-conserved in human Lats (Lys-696) and Ndr (Lys-80) kinases at a similar position in the NTR – Mob1 structures solved to date (Figure 2B and C) (44,45). This would position a positively charged side chain above the key HM phosphorylation site (Cbk1 Thr-743, Ndr1 Thr-444, Lats1 Thr-1079) with additional conserved salt bridges forming between Cbk1 Glu-336 and Arg-746 (Figure 1B). In Lats, Glu-689 is crucial for interaction with Mob1 in humans (44); it is equivalent to Glu-336 of Cbk1 and otherwise is extremely highly conserved (Figure 2C). Overall, the revised Cbk1-Mob2 structure supports the presence of a robust, coordinated set of electrostatic linkages among the Mob cofactor, kinase HM, and kinase NTR region. These form a likely regulatory mechanism distinctive to Ndr/Lats kinase – Mob cofactor complexes that are core elements of hippo pathways across the eukaryotic tree.

### The Dbf2-Mob1 structure exhibits structural similarity to Cbk1-Mob2

Budding yeast Mob1 is among the most extensively biochemically characterized Mob coactivators, with a significant number of its properties elucidated using *mob1* temperature sensitive alleles discovered in genetic screens (11,13,17). However, Mob1’s crucial interaction interface with Dbf2 is not well characterized. To more completely understand this complex and understand its similarities and differences with Cbk1 and Lats1 we solved the structure of Mob1^79-314^ bound to Dbf2^NTR^ (Dbf2^85-173^) to 3.5 Å resolution (Figure 4A). Like Cbk1^NTR^-Mob2, the Dbf2 NTR region forms a two-helix hairpin, with αMob helices that engage in binding interactions with each other and with the Mob1 cofactor. Many of these contacts are similar to those found in Cbk1^NTR^-Mob2 (Figure 4B, compare with Figure 1B).

**Figure 4.**
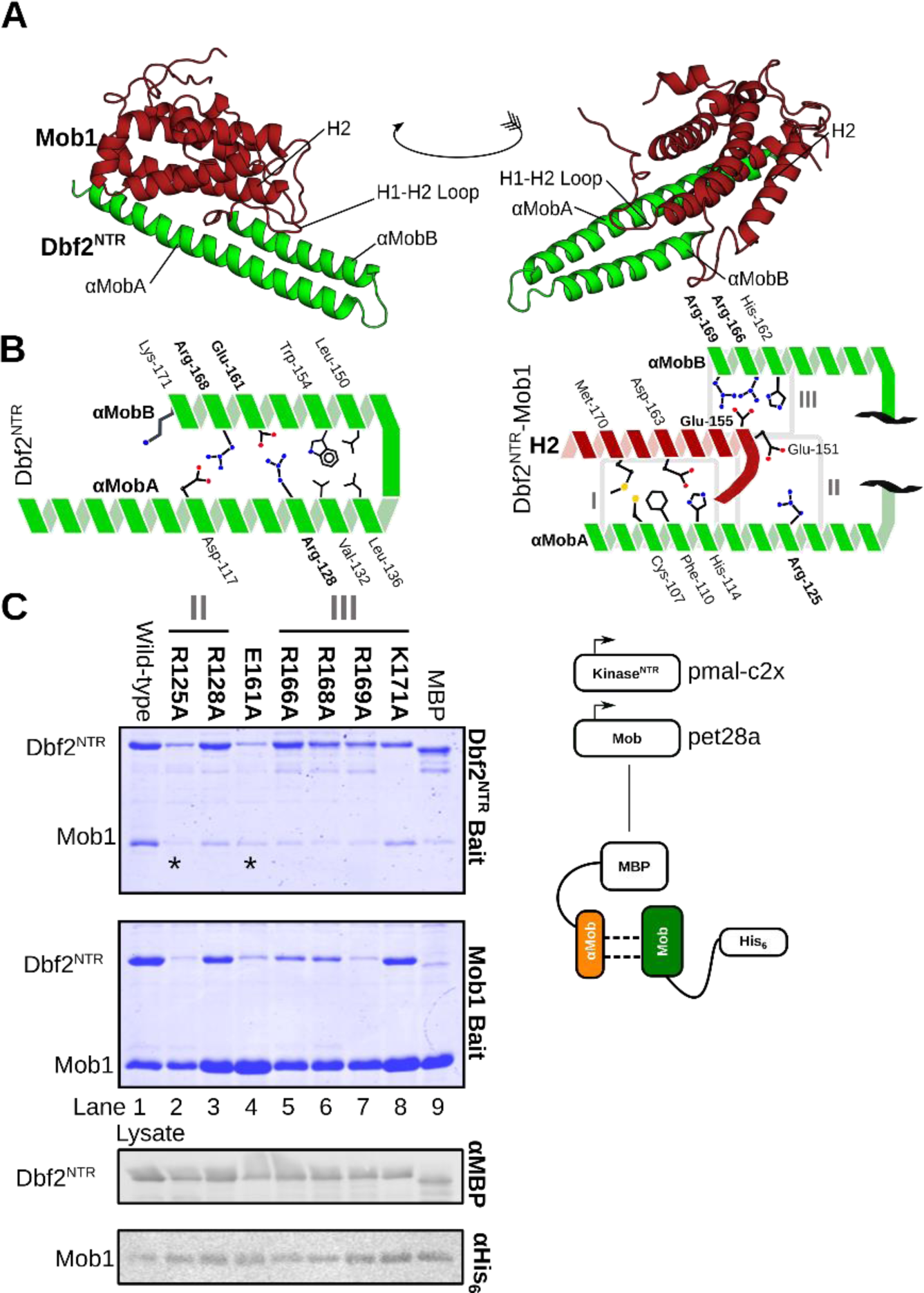
Dbf2 interacts with Mob1 through conserved residues. (A) The crystal structure of Dbf2 ^NTR^ -Mob1 with conserved motifs labeled. Dbf2, green; Mob1, dark red. (B) Interaction architecture of interacting residues within the Dbf2^NTR^ helices αMobA and B (left) and between Dbf2^NTR^ and Mob1 (right). Conserved residues are shown in bold. (C) BL21(DE3) RIL co-expressing MBP-fused Dbf2^NTR^ (85–173) and His6-fused Mob1 (79–314) followed by co-purification of the cognate binding proteins with amylose (top) and nickel (middle) resin. We separated the complexes using SDS-PAGE and stained with Coomassie. Lower bottom panels represent Western blots of the lysate as a loading control.

We assessed the characteristics of this complex by performing interaction assays with point mutants in conserved residues identified in the crystal structures. We found that fusion to maltose binding protein (MBP) dampens the aggregation of unstable Ndr/Lats-Mob complexes and permits detection of interactions when co-expressed with Mob in a bacterial system. Pull downs were performed in reciprocal with either MBP-Dbf2^NTR^ (Figure 4C, top) or Mob1-His6 (Figure 4C, middle) with input controls shown (Figure 4C, bottom). MBP-Dbf2^NTR^ Mob1 interaction assays demonstrated that Dbf2 Arg-125, Arg-166, and Arg-169 contribute strongly to the interaction, as predicted by the crystal structure (Figure 4B and Figure 4C lanes 2, 5, and 7). Mutation of non-Mob-binding Arg-128, Glu-161, and Arg-168 either ablated or abolished the interaction (Figure 4B and Figure 4C lane 3, 4 and 6), indicating integrity of the NTR is crucial to formation of the complex similar to the intra-αMob interaction between Lats1 Arg-660 and Glu-689 (45). Interestingly, Lys-171 (Figure 4C, lane 8) participates minimally in Mob1 binding despite strong charge conservation across the eukaryotic world. Thus, conserved residues in the Dbf2-Mob1 structure participate in binding.

### Conserved regions cohere Ndr/Lats Mob complexes

To gain further insight into the conserved structural mechanisms that might mediate the formation of the binding interface, we compared the structure of Cbk1^NTR^-Mob2 and Dbf2^NTR^-Mob1 in conserved interface II and III regions (Figure 5). Overall, we found both complexes to be conformationally similar to previously published structures of Mob1/Lats1 (44,45) and Mob1/Ndr2 (43). Both Cbk1^NTR^ and Dbf2^NTR^ form a two-helix bundle (αMobA and αMobB) in which the helix 2 (H2) of the Mob protein incorporates into a cleft formed between the two αMob helices (Figure 5A). We also examined a highly conserved interface (interface II) that holds the αMobA and αMobB helices together (Figure 5A see also Supplementary Figure 2). Similar to Lats-Mob1, Dbf2 Arg-125 and Arg-128 protrude from αMobA to hold Glu-161 on αMobB (Figure 5A, left panel) akin to interface II in the Mob1-Lats1^NTR^ structure (Figure 5A, center panel) (45). Dbf2 Arg-125 interacts with Mob1 Glu-151, which is conserved in Mob1 co-activator subfamily (17), and with Dbf2 αMobB His-162 (Figure 5A, left panel). Similarly, the Cbk1-Mob2 structure reveals a salt bridge between Arg-307 and Glu-336 (Figure 5A, right panel).

**Figure 5.**
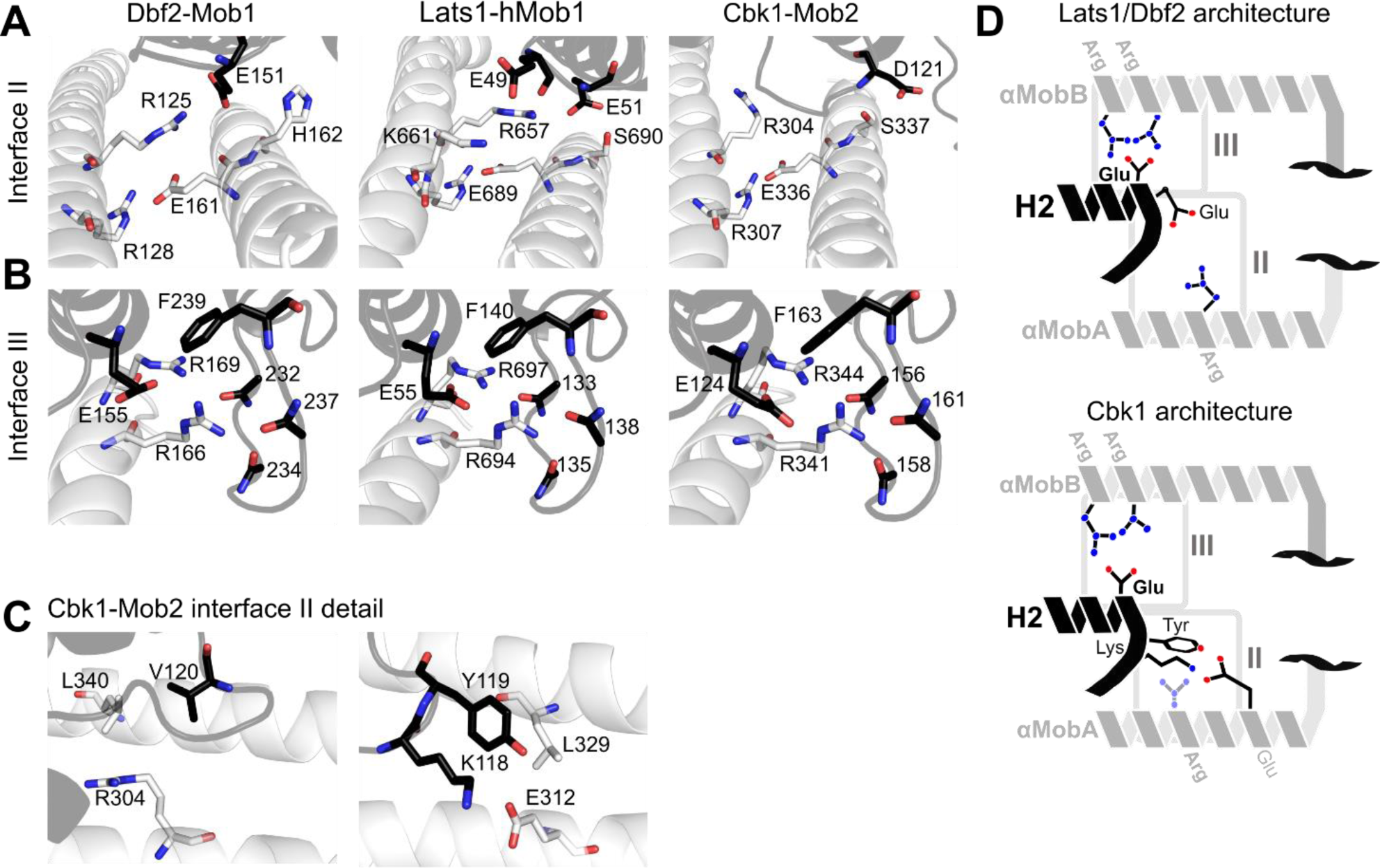
Conserved regions cohere Ndr/Lats ^NTR^-Mob complexes. Mob is shown in black, Ndr/Lats kinase ^NTR^ is shown in grey. (A) Comparison of interface II from Lats1 ^NTR^-hMob1 (PDB code 5B5W, center panel) with Cbk1 ^NTR^-Mob2 (right panel) and Dbf2 ^NTR^-Mob1 (left panel). Highlighted residues shown as sticks are important for the interaction. (B) Comparisons at interface III with Lats1 ^NTR^-hMob1 (center panel), Cbk1 ^NTR^-Mob2 (right panel) and Dbf2 ^NTR^-Mob1 (left panel). (C) More detailed view at interface II of the Cbk1 ^NTR^-Mob2 complex showing the orientation of Cbk1 Arg-304 in relation to Leu-340 and Mob2 Val-120 (on the left) and Cbk1 Glu-312 and Mob2 Lys-118, Tyr-119 (on the right). The interaction between Lys-118/Tyr-119 of Mob2 and Glu-312/Leu-329 of Cbk1 is distinct from Lats and Dbf2-Mob1 complexes. (D) Generalized architecture of Dbf2/Lats1-Mob1 in relation to Cbk1-Mob2. Conserved binding interfaces II and II as shown in (A-C) are boxed. Basic residues, blue; acidic residues, red.

Similar to Mob1/Lats1 (45), several highly conserved basic residues form an interface (interface III) to stabilize the Mob-NTR interaction (Figure 5B). Dbf2 Arg-166 and Arg-169 contact Mob1 Glu-155 (Figure 5B, left panel). It was shown that Lats/Mob1 shows a similar organization with the conserved Lats Arg-694 and Arg-697 interacting with Mob1 Glu-55 (Figure 5B, center panel) (45). Similarly, Cbk1 Arg-341 and Arg-344 form an electrostatic interaction with Mob2 Glu-124 (Figure 5B, right panel). Taken together, these data demonstrate how the geometry of the Mob1-NTR interaction at interface III is strongly conserved; Lats1, Dbf2 and Cbk1 bind their cognate Mobs in a nearly identical fashion, especially in interface III, with some differences in interface II described below.

While Cbk1 Arg-307 bridges with Glu-336 on αMobB similar to the inter-helical cohesion between Lats1 Arg-660 and Glu-689, Cbk1 Arg-304 does not interact with Mob in the crystal structure as Lats1 Arg-657 and Dbf2 Arg-125 do (Figure 5A), instead being sterically pushed towards the center of Mob2 by the hydrophobic Val-120 and Cbk1 Leu-340 (Figure 5C, left panel). Replacing this interaction, Mob2 Lys-118 and Tyr-119 extend from the H1-H2 loop and contact Cbk1 Glu-312, forming a salt bridge and a hydrogen bond, respectively, and laying in a pocket formed at the intersection of the two αMob helices (Figure 5C, right panel). Glu-312 is not conserved in Dbf2, but found in other known Ndr/Lats-subfamily kinases in metazoans. Thus, Cbk’s interaction with Mob2 partially occurs through an interaction distinct from Lats1/Dbf2 (Figure 5D).

### Budding yeast Mob1 and Mob2 have intrinsic binding specificity for their cognate kinases

Ndr and Lats kinases, while highly similar, are recognizable as distinct AGC kinase subfamilies in eukaryotes separated by over a billion years of evolution. Moreover, they largely act in non-overlapping hippo pathway signaling systems with different physiological roles. The remarkable functional segregation of the highly similar Ndr and Lats kinases may be reinforced by association of their NTR regions with specific Mob-family coactivators that are distinct from one another over large evolutionary distances. While formation of specific kinase-coactivator complexes can be achieved by partitioning proteins in space, time, or cell type, direct biochemical evidence indicates that Mob coactivators have intrinsic binding specificity for the Ndr or Lats NTR regions. In *S. pombe*, for example, the Lats-related kinase Sid2 binds Mob1 and the Ndr-related kinase Orb6 binds Mob2, forming “cognate” Sid2-Mob1 and Orb6-Mob2 complexes without “non-cognate” kinase-coactivator cross-interaction (39,49). Genetic evidence supports a similar organization in *S. cerevisiae*, but this has not been demonstrated biochemically. We therefore examined the association specificity of Dbf2 and Cbk1 with Mob1 and Mob2.

Unfortunately, conventional binding assays are highly problematic for *in vitro* assessment of association of Ndr/Lats kinases with Mob coactivators. NTR regions of Ndr/Lats kinases behave exceptionally poorly in expression and purification. We find that Cbk1^NTR^ and Dbf2^NTR^ form insoluble and soluble aggregates, making them essentially intractable for meaningful solution binding assays. This is likely broadly true for Ndr/Lats NTR regions: notably, Lats(NTR)-Mob1 complexes used in recent crystallographic studies were produced by mixing chemically solubilized, denatured Lats(NTR) with purified Mob1 (REF). We also find that Cbk1 and Dbf2 kinase constructs containing the kinase NTR region express and purify poorly, and that these problems are remedied by co-expression of the native Mob cofactor. Thus, we used co-expression to evaluate binding and specificity of NTR region – Mob binding.

To test NTR region – Mob cofactor binding specificity we co-expressed MBP-fused Dbf2^NTR^ or Cbk1^NTR^ with either His_6_-fused Mob1^79-314^ or zinc-binding His_6_-Mob2^45-278^ using a two plasmid system in which both components are independently expressed in bacterial cells to facilitate a stable interaction between the proteins. We purified His_6_-Mob with Ni-NTA resin and assessed copurification of kinase NTR by SDS-PAGE and Coomassie stain. These data show copurification of Dbf2^NTR^ with Mob1 and Cbk1^NTR^ with Mob2, indicating that cognate NTR-Mob complexes formed under two plasmid co-expression conditions (Figure 6A, lanes 1, 4). There is no evidence that non-cognate Dbf2-Mob2 formed (Figure 6A, lane 3). Consistent with analysis of *S. pombe* orthologs(49), formation of very small amounts of Cbk1^NTR^ -Mob1 is evident (Figure 6A, lane 2). We also performed bi-cistronic expression of kinase His_6_-NTR and GST-Mob, which at least in principle better coordinates translation and stoichiometry of the two components. This provided a more sensitive assessment of component binding capability. We found robust expression and co-purification of cognate kinase NTR – coactivator pairs: Dbf2^NTR^ with Mob1 and Cbk1^NTR^ with Mob2 (Figure 6B). Strikingly, the proteins appeared to be highly unstable when expressed as non-cognate pairs, with no indication that Dbf2^NTR^-Mob2 or Cbk1^NTR^ - Mob1 complexes formed (Figure 6B). We also assessed interaction of these proteins using yeast two-hybrid analysis, and saw consistent results with this orthogonal assay (Supplementary Figure 3). Overall, our data indicate that that limited cross-interaction occurs between the non-cognate complexes suggesting Dbf2-Mob1 and Cbk1-Mob2 act as insulated complexes.

**Figure 6.**
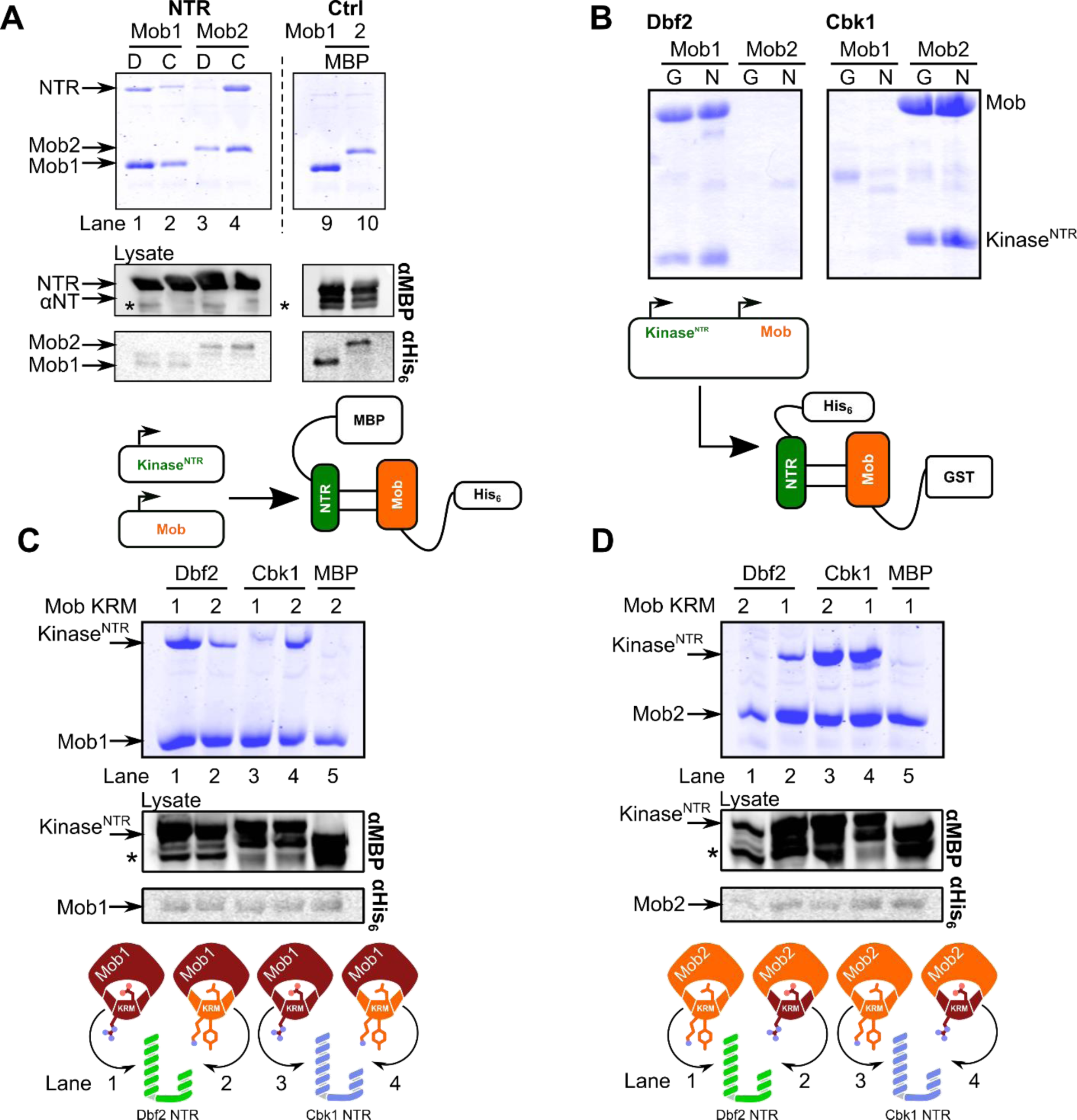
Ndr/Lats-Mob binding is specific and determined by multiple regions. (A) Dbf2 and Cbk1 form specific complexes with Mob1 and Mob2 *in vitro*. MBP-fused Dbf2 ^NTR^ (85–173), labeled as “D”, and Cbk1 ^NTR^ (251–351), labeled as “C”, were co-expressed with His6-fused Mob1 (79–314) and Mob2 ^V148C Y153C^ (45–278) in BL21(DE3), isolated with nickel chromatography, separated with SDS-PAGE, and stained with Coomassie (lanes 1-4). Lanes 9 and 10 are MBP controls. The dotted line demarcates a division in the gel; see Supplementary Figure 4 for full gel. (B) Complexes of Dbf2 ^NTR^ and Cbk1 NTR with Mob1 and Mob2 were expressed complexes from a single bi-cistronic plasmid in BL21(DE3) and isolated by glutathione (G) and nickel (N) resin. (C) The Mob1 and (D) Mob2 kinase restrictor motif (KRM) interacts with Ndr/Lats ^NTR^ and is important for specificity. (C) Mob1 containing either the wild-type (1) or Mob2 (2) KRM and Dbf2 ^NTR^ or Cbk1 ^NTR^ were co-expressed in BL21(DE3) and purified with nickel resin using Mob as a bait. Lane 1, Dbf2 ^NTR^ – Mob1^WT^; 2, Dbf2 ^NTR^ -Mob1 ^Mob2KRM^; 3, Cbk1 ^NTR^ -Mob1 ^WT^; 4, Cbk1 ^NTR^ –Mob1^Mob2KRM^. (D) Mob2 ^V148C Y153C^ containing the Mob1 (1) or Mob2 (2) KRMs co-expressed with Dbf2 ^NTR^ or Cbk1 ^NTR^ similarly. Lane 1, Dbf2 ^NTR^ -Mob2 ^WT^; 2, Dbf2 ^NTR^ -Mob2 ^Mob1KRM^; 3, Cbk1 ^NTR^ -Mob2 ^WT^; 4, Cbk1 ^NTR^ -Mob2 ^Mob1KRM^. Lane 5 in all cases is a control with either Mob1 ^Mob2KRM^ or Mob2 ^Mob1KRM^. The cartoon (bottom) describes the experiment; Mob1, dark red; Mob2, orange. Mutants: Mob1 ^Mob2KRM^, Mob1^R149K G150Y E151V^; Mob2 ^Mob1KRM^, Mob2^K118R Y119G V128E^. (*) Denotes degradation products of the NTR.

The Cbk1^NTR^-Mob2 crystal structure revealed that a non-conserved strand of Cbk1^259-277^, required for expression of the full-length Cbk1-Mob2 complex, formed contacts with the back of Mob2 (Supplementary Figure 4A). Preliminary mutational analysis of the Cbk1^NTR^ in interfaces II and III failed to abolish binding to Mob2, further suggesting an alternative binding site (data not shown). Indeed, truncation of Cbk1 to a minimal construct Cbk1^251-303^ (Cbk1 ^αNT^), containing this strand and first 27 amino acids of the αMobA helix, was sufficient to bind zinc-binding Mob2 with wild-type affinity despite the lack of the conserved electrostatic interactions found in Dbf2/Lats interfaces II and III (Supplementary Figure 4B, lane 5-8). Furthermore, Cbk1 ^αNT^ expressed with Mob2 lacking the zinc binding mutation is sufficient to express the coactivator solubly, suggesting a novel Cbk1-Mob2 interaction motif replaces the metal binding site (data not shown). In contrast, Dbf2^85-124^(Dbf2 ^αNT^) cannot bind Mob1 or Mob2 and is rapidly degraded, suggesting the bi-helical motif is required for stabilizing the Dbf2 NTR (Supplementary Figure 4B, lanes 5, 7).

### The Mob kinase restrictor motif (KRM) constrains kinase binding to a single cofactor

All crystallized Ndr/Lats kinases interact with a conserved loop between helices 1 and 2 (H1 and H2) of the associated Mob protein (Supplementary Figure 5) (17,44,45). Alignment of this H1-H2 region in budding yeast and human Mob proteins identifies a highly conserved (L,P)XXX(D,N) motif (Supplementary Figure 6). In budding yeast Mob1 the conserved and essential proline Pro-148 appears to stabilize the H1-H2 loop (17), and the solvent-exposed asparagine Asp-152 initiates helix 2. Intriguingly, this motif in Mob H1-H2 also separates Mob1 and Mob2 subfamilies: the three variable amino acids between these positions are subfamily-specific. Mob1-family proteins are characterized by (L,P)XGE(D,N) sequences, while Mob2 proteins lack strong conservation of variable H1-H2 positions (Supplementary Figure 6). In budding yeast, the Mob1 H1-H2 tripeptide sequence is RGE, while in Mob2 the corresponding tripeptide sequence is KYV. We surmised that the H1-H2 loop tripeptide contributes to NTR region-Mob cofactor binding specificity. As discussed further below, we refer to this H1-H2 tripeptide as the “Kinase Restrictor Motif” or “KRM”.

To assess the H1-H2 region’s role in kinase - Mob binding specificity we generated mutant Mob1 and Mob2 proteins with H1-H2 “KRM” sequences of interest swapped. Specifically, we produced Mob1 with the R149K G150Y E151V alteration and Mob2 with the K118R Y119G V120E alteration. For brevity we refer to these mutant constructs as Mob1^Mob2KRM^ and Mob2^Mob1KRM^. We co-expressed these constructs with maltose binding protein fusions of Dbf2^NTR^ and Cbk1^NTR^. As shown in Figure 6C, the Mob1^Mob2KRM^ mutant protein formed complexes with Dbf2^NTR^ less well than wild type Mob1, and associated much more dramatically with the non-cognate Cbk1^NTR^ construct than wild-type Mob1 (Figure 6C). Similarly, we found that the Mob2^Mob1KRM^ mutant protein robustly formed complexes with Dbf2^NTR^. Notably, while there is some experimental evidence that direct Ndr-Mob1 interaction may occur, there is to date negative evidence that Mob2 can interact with Lats kinases (Figure 6A and B) (8,49–51). Since the H1-H2 tripeptide is a feature different in Mob1 and Mob2 families that confers or enhances their ability to bind non-cognate Ndr/Lats kinases, we henceforth label it a “Kinase Restrictor Motif” or “KRM”.

We next determined if the interaction of Mob KRM mutants with the Ndr/Lats^NTR^ were consistent with the established Ndr/Lats-Mob binding architectures and not artifactual or otherwise due to aggregation or nonspecific interactions within the complex. Mob1 ^Mob2KRM^ and zinc-binding Mob2 ^Mob1KRM^ were expressed with MBP-fused Dbf2 and Cbk1 mutants with mutations in conserved residues in interfaces II and III (Supplementary Figure 7). Alanine substitutions for Dbf2 Arg-125 and Arg-128 ablated binding to chimeric Mob2 ^Mob1KRM^, similar to that observed in Mob1 and consistent with the Dbf2-Mob1 and Lats1-hMob1 crystal structures (Supplementary Figure 7A, lanes 4-5) (44,45). Importantly, substitution of Cbk1 Arg-304 for alanine does not affect binding of Cbk1 to Mob1 ^Mob2KRM^, consistent with this residue’s lack of Mob interaction and demonstrating Cbk1’s binding is through a Mob2-like binding interface (Figure 5C, and Supplementary Figure 7B, lane 4). Mutations in interface III’s conserved Dbf2/Cbk1 Arg-166/341 and Arg-169/344 residues, found in all known Lats/Ndr kinases, further ablates the interaction (Supplementary Figure 7A,B, lanes 7-9). These findings suggest the Mob KRM interacts through the NTR in a manner consistent with its placement in the crystal structure.

## Discussion

### Ndr/Lats kinase activation: the role of HM phosphorylation and Mob binding

Phosphorylation at the hydrophobic motif (HM) on Thr-743 by an upstream kinase activates Cbk1 and the C-terminal activation loop (AL) region harbors an auto-phosphorylation site (Ser-570) which is required for full activation. Thr-743 found near interface III of the Cbk1^NTR^ is positioned close to the highly conserved basic residues in that region such as Arg-343 (Figure 2). We further note that the position of this residue is possibly fixed by the Mob2 interaction, and may replace the basic binding pocket found in most AGC kinases. We suggest that this region may be in common in Ndr/Lats kinases due to the high similarity between the Cbk1-Mob2 structure and previously published complexes of Lats1^NTR^-Mob1 as well as our Dbf2^NTR^-Mob1 structure (Figure 4) (44,45). Furthermore, amino acid conservation in this region is highly suggestive that this may reflect a general mechanism for activation of the Ndr-family of kinases (Figure 2B and C).

In order to gain insight into how phosphorylation at these two regulatory sites may promote kinase activity, we performed MD simulations on the complex to address the possible allosteric role of AL and particularly HM phosphorylation (Supplementary Figure 8); as the new Cbk1^NTR^-Mob2 crystal structure necessitated the revision of the activation model that we had formerly put forward (46). For MD simulations a hybrid model was constructed using the new, corrected Cbk1-Mob2 crystal structure and PKB/Akt structures (48). In contrast to our earlier findings, where we relied on a less complete Cbk1-Mob2 crystal structure to construct an MD starting model, the new simulations did not indicate long range movements at αMobB (previously referred to as N-linker) between αMobA and the AGC kinase core. In the light of the corrected model, however, this is now not surprising. In the old model the HM region could only be partially traced and the amino acid register of the αMobA helix was misinterpreted. In the revised crystallographic model, the full HM region became visible and the register at the N-terminus was corrected.

Although the new MD simulations indicated some small-range differences regarding the movements of critical Cbk1^NTR^ arginine residues directly contacting the phosphorylated side chain of Thr-743 (e.g. Arg-343, located at interface III) in the conformational dynamics of the non-phosphorylated and phosphorylated complexes, this computational approach failed to reveal the exact mechanistic role of Ser-570 and Thr-573 phosphorylation in Cbk1 activation: (1) biochemically feasible modeling of the critical αC segment became possible only if main-chain restraints were applied, (2) trajectories even for long (∼1 μM) simulations were too sensitive to small differences in the starting models as the outcome varied depending on what parts of the models were allowed to move freely or restraint, or if the models contained ATP or Mg-ATP for example (Supplementary Figure 8). Despite of this, the Cbk1 HM region may promote the ordering of αC in the phosphorylated complex and thus influences activity through this kinase region known to be frequently involved in allosteric regulation. Overall, our structural and MD analysis is consistent with a mechanistic model where Ser-570 phosphorylation at the activation loop and particularly Thr-743 phosphorylation at the HM promote the optimal positioning of the otherwise flexible αC. This mechanism clearly resembles to how HM phosphorylation and binding at the so-called PIF pocket affects the activity of other better known AGC kinases (48) (Figure 3).

### Evolutionarily conserved features of the Ndr/Lats kinase-coactivator interface

In humans, Lats kinases restrain cell proliferation by keeping the YAP/TAZ transcription factors out of the nucleus. Ablation of signaling within the pathway, such as reductions in Ndr/Lats catalytic activity or 14-3-3 levels, which promote YAP/TAZ cytosolic sequestration can cause cell overgrowth and transformation (52). Our findings identify conserved basic amino acids in the NTRs of Ndr/Lats kinases that are probably essential for their functionally critical interaction with Mob cofactors. Notably, mutations affecting these basic amino acids in human Lats1 and 2 have been linked to cancers: R657C, R694C, and R697G mutations in Lats1 strongly correlate with uterine, ovarian, and papillary renal cell cancers, and Lats2 R623W correlates with melanomas (53). We suggest that these Lats1 and 2 alleles have weakened Mob1 binding, disrupting cofactor interaction and the complex’s function.

Dbf2 and Cbk1 are conserved AGC kinases in the Ndr/Lats family and are evolutionarily related to their human counterparts. The conserved arginines on the αMobA and B helices of Dbf2, Cbk1, Lats and Ndr contain a conserved RxxR motif (see Figure 1D). Dbf2/Lats uses this motif to bind its Mob factor, whereas Cbk1 does not appear to. While the second arginine appears to electrostatically cohere the αMob helices, the function of Cbk1’s Arg-304 remains unknown, as it does not bind Mob2. However, Cbk1 Arg-304 may have some role in stabilizing the HM binding site in the kinase-coactivator complex as Cbk1^HM^ Arg-746 interacts electrostatically with the conserved Glu-336 which is coordinated by the RxxR of Cbk1^NTR^ (Figure 2B and C). Mob1 binds Dbf2 RxxR (involving Arg-125 and Arg-128) directly to form interface II (Figure 4B), suggesting roles for the cofactor in kinase regulation through modulation of the HM binding site if the structures are homologous.

### What is the specificity of Ndr kinases for the Mob cofactor?

Human Ndr kinases have been noted to interact with Mob1 (40), an interaction that may depend on phosphorylation of Mob1 Thr-74 (54,55). Ser-174 in *S. cerevisiae* is homologous with this residue, though its phosphorylation state is unknown. This non-cognate interaction has not been observed in *S. pombe* or *S. cerevisiae* model systems in previous *in vitro* assays. We find that while Cbk1 strongly binds Mob2 *in vitro*, the kinase’s interaction with Mob1 interaction is weak and unstable. Our yeast two-hybrid data suggests that Cbk1-Mob1 association can occur and previously published high throughput studies have obtained similar results (50), however the physiological relevance of this interaction is not clear. We note that isothermal titration calorimetry (ITC) data for the human Ndr^NTR^-Mob1 interaction has been obtained, however, the oligomeric state of the Ndr^NTR^ used was not specified (43). In our hands, we find Dbf2 and Cbk1 NTRs form aggregates when purified without their associated Mob, a result also observed with human Lats^NTR^ (44,45). Therefore, we suggest additional biochemistry should be performed to determine the exact interaction state between these Ndr-Mob complexes.

In immunoprecipitation experiments from cultured cells, mutations in interfaces II and III of human Ndr1^NTR^ abrogate its association with Mob1, but not of Mob2, suggesting binding to Ndr1 occurs through distinct mechanisms (51). Whether the observed interaction is direct or through an adapter has not been determined, though point mutations in the NTR suggest binding uses homologous residues to Lats1-hMob1 (7,8,17,45,56). Human Mob1 has been shown to interact with Mob2 (57), an interaction also observed in *S. cerevisiae* (35), suggesting a potential oligomerization mechanism for the cofactors. Further research will be required to fully elucidate the nuances of these interactions.

The ability of human Mob2 to “out-compete” Mob1 for binding to Ndr1 as reported by Kohler et al. is consistent with the observed strength of the Cbk1-Mob2 interaction (51). No mutation in Ndr^NTR^ is capable of abrogating binding to Mob2 (51), a phenomenon observed with Cbk1 in the hands of the authors as well. Mob1 localizes to spindle pole bodies, but not the bud neck, in the absence of Dbf2 (12); the non-Dbf2-binding *Mob1-77* mutant has a similar effect (9). In contrast, Mob2 in the absence of Cbk1 displays a completely diffuse localization pattern consistent with degradation in the absence of the stabilizing Cbk1^NTR^ (26,32); this is likely due to the strand of Cbk1 ^αNT^ (Cbk1^259-277^) stabilizing Mob2 (Supplementary Figure 4). As this strand appears to not exist in Ndr, it may be difficult to draw parallels between human and *S.cerevisiae* biochemistry within the Ndr/Lats paradigm.

### The Kinase Restrictor Motif of Mob plays a key role in defining the specificity of kinase-coactivator interactions

Alignment of Mob cofactor protein sequences display subfamily-specific sequences which might divide Mob1 from Mob2 and thus impart specificity on kinase binding (Supplementary Figure 6). The three amino acids identified as the KRM extend from the N-terminus of Mob helix I and orient towards the predicted kinase interaction surface in Mob structures solved previously (17,35,58). In Ndr/Lats^NTR^- Mob, the KRM lays into a binding cleft in the NTR (Supplementary Figure 5). Many inactivating mutations in *S. cerevisiae* and human Mob1 map to this motif or residues surrounding it (13,17,45).

Comparing the KRMs of eukaryotic Mob1 and 2 subfamilies may reveal similarities in cofactor function. Mob1’s high conservation in this region is consistent with its generally conserved role in binding Lats subfamily kinases and in cell cycle exit. Mob2 is more diverse, potentially reflecting the larger variety of functions performed by the Ndr subfamily of kinases in eukaryotic systems. Human Mob2 may interact with Ndr through Arg-48 and Glu-49 of its KRM. In contrast, *Drosophila* Mob2, which contains Ala/Gly in the place of Arg/Glu, may not utilize the motif to bind Tricornered. Such large differences within the normally conserved Ndr-Mob families may explain the difficulty of abolishing the human Ndr1-Mob2 interaction through point mutations at conserved residues in previous work (51).

In our revised structure the conserved arginines in Cbk1’s Mob2-binding NTR (RxxR) associate with Arg-746 in the kinase’s C-terminal regulatory hydrophobic motif (HM) (Figure 2). The position of Mob2^KRM^ directly covering the highly conserved HM binding site in Cbk1 suggests a role for regulation. As hMob1/Mob1^KRM^ directly binds the homologous domain through Glu-51/151, we hypothesize Mob1 modulates the conformation of the putative Lats/Dbf2 HM binding region and subsequently controls the kinase.

Our analysis here indicates that the Ndr/Lats^NTR^-Mob interface is a common structural platform through which kinase-cofactor binding is mediated, however, amino acid variations in key positions contribute to subgroup and organism-specific differences. While the first Mob crystal structures suggested that a conserved and acidic surface was responsible for interactions with Ndr/Lats through the kinase N-terminus, the reality may be more complex. Some Mob cofactors may bind and activate their cognate kinases through a conserved set of charged residues; others may bind using distal regions and use these residues for activation only, or for alternative functions of Mob. More research is required to elucidate the mechanisms of binding, specificity, and activation of these unique complexes and their role in physiology.

## Methods

As reported previously (17), we were unable to stably express and purify monomeric Mob2 in an *E.coli* overexpression system. Therefore, to produce suitable protein for crystallization and biochemistry, we engineered Mob2 ^V148C Y153C^ which recapitulates a zinc-binding motif found in most metazoan Mob2 orthologs as well as in *S. cerevisiae* Mob1. Unless otherwise noted, interaction data and the Cbk1^NTR^- Mob2 crystal structure were obtained using this mutant.

### Expression and Purification of Proteins

To purify the zinc-binding Cbk1^NTR^-Mob2 ^V148C Y153C^ complex, a modified bi-cistronic pBH4 vector containing His_6_-Cbk1 (Cbk1 ^NTR^, residues 251-351) and zinc-binding GST-Mob2 ^V148C Y153C^ (residues 45-287) expressed from a single T7 promoter was cloned by ligating Cbk1 residues 251-756 and Mob2 45-287 into the backbone, then mutating Cbk1 Phe-352 to a stop codon using the QuickChange technique (Agilent). Val-148 and Tyr-153 were mutated to cysteine using the same technique. Protein was overexpressed in BL21(DE3) RIL (Agilent) at 37 °C in Terrific Broth (TB) containing 40 μM zinc chloride. After growth to mid-log phase, IPTG was added to 0.2mM and expressed overnight at 37 °C. Cell pellets were lysed by sonication, purified using nickel chromatography, and tags cleaved using Tobacco Etch Virus (TEV) protease overnight on wet ice. Cleaved protein was further purified using a hand-poured SP-Sepharose (GE) column and eluted with a gradient from 50-2000 mM NaCl, then buffer exchanged into 20 mM HEPES pH 7.4, 300 mM NaCl and concentrated to 55 mg/ml.

To purify the Dbf2 ^NTR^ -Mob1 complex, a bi-cistronic vector containing His_6_-Dbf2 ^NTR^ (85–173) and untagged Mob1 (79–314) expressed from a single T7 promoter was cloned and overexpressed in BL21 (DE3). Mob1 (79–314) contained the N-terminal amino acid sequence ENLYFQGS as a byproduct of cloning. TB cultures containing 40 μM zinc chloride were inoculated, grown to mid-log phase, induced with IPTG to 0.2 mM, and expressed overnight at 24 °C. Cell pellets were lysed by sonication. Protein was purified using nickel chromatography and cleaved overnight on wet ice using TEV protease. Residual TEV and nickel-binding contaminants were reabsorbed using nickel resin, final protein was then buffer exchanged into 100 mM sodium phosphate pH 7.4, 50 mM KCl, 100 mM L-arginine HCl, and concentrated to 30 mg/ml. L-arginine was present throughout the purification to prevent oligomerization of the complex and did not interfere with nickel chromatography.

### Crystallization of Cbk1-Mob2, Cbk1 ^NTR^ -Mob2, and Dbf2 ^NTR^ -Mob1 complexes

Ndr/Lats ^NTR^ -Mob crystals were obtained at 22.3 °C. Cbk1 ^NTR^ -Mob2 was crystallized using the hanging-drop method; well solution containing 500 μL 100mM MES/Imidazole pH 6.5, 30 mM CaC_2_, 30 mM MgCl_2_, 12.5% (w/v) PEG1000, 12.5% (w/v) PEG3350, 12.5% (v/v) MPD produced crystals. Dbf2 ^NTR^ -Mob1 was crystallized using the microbatch method under 200 μL mineral oil (Fisher BioReagents); crystals were obtained in 100 mM Bicine/Tris pH 8.5, 30 mM LiCl, 30 mM NaCl, 30 mM KCl, 30 mM RbCl, 12.5% (w/v) PEG1000, 12.5% (w/v) PEG3350, 12.5% (v/v) MPD. All crystals were frozen in liquid nitrogen without the addition of further cryoprotectant. Cbk1 ^NTR^ -Mob2 and Dbf2 ^NTR^ - Mob1 diffracted to 2.80 Å and 3.50 Å, respectively.

Crystallization of the Cbk1-Mob2-Ssd1 complex was identical as previously published (46). To increase the redundancy of our crystallographic data and to maximize the gained experimental information, we merged two independent datasets collected on this complex (Table 1). This helped to increase the resolution range to 3.15 Å, meaning an additional ∼15% experimental data on top of duplicated multiplicity. Data were collected at 100 K on the PXIII beamlines of the Swiss Light Source (Villigen, Switzerland) and at the Advanced Photon Source (Argonne, IL). Data was processed with XDS (doi: 10.1107/S0907444909047337) and the structures of Cbk1 ^NTR^ -Mob2, and Dbf2 ^NTR^ -Mob1 complexes were solved by molecular replacement with PHASER (doi: 10.1107/S0021889807021206). The MR search identified single Mob-αMob complexes with a searching model of Lats1-hMob1 (PDB ID: 5BRK). Structure refinement was carried out using PHENIX (doi:10.1107/S0907444909052925) and structure remodeling and building was performed in Coot (doi: 10.1107/S0907444910007493).

### Pull-down experiments

Dbf2 (85–173) and Cbk1 (251–352) were cloned into pMAL-c2x (New England Biolabs) and fused to an N-terminal MBP tag. *S.cerevisiae* Mob1 (79–314) and zinc-binding Mob2 ^V148C Y153C^ (45–278) were cloned into pET28a (EMD Biosciences). Plasmids were mutated using the QuickChange technique (Agilent). Constructs were co-transformed into BL21(DE3) or BL21(DE3) RIL, grown in TB at 37 °C for two hours, induced with IPTG to 2 mM, and grown overnight at 18 °C or 24 °C. Cell pellets were lysed by sonication and raw protein measured by Bio-Rad Protein Assay (Bio-Rad). Lysates were normalized, incubated with nickel resin (Qiagen) or amylose resin (New England Biolabs), washed twice with 20 mM imidazole, and eluted in 300 mM imidazole (nickel) or by boiling resin in 1X SDS-PAGE loading dye (amylose). Purified protein was separated using 15% SDS-PAGE gels and visualized using an Odyssey infrared imager (LI-COR Biosciences).

### Immunoblotting

Bacterial cell lysates normalized in raw protein concentration were separated on 15% SDS-PAGE gels and transferred to polyvinylidene difluoride (PVDF) membranes using Dunn’s modified transfer buffer (59). Blots were blocked using Odyssey Blocking Buffer (LI-COR Biosciences) for 30 minutes and probed with mouse αMBP or rabbit αHis_6_ antibodies for 1 hour at room temperature. Membranes were washed three times with TBS-T, probed with fluorophore-conjugated IRDye 680LT goat αMouse or IRDye 800CW goat αRabbit (LI-COR Biosciences) for 30 minutes, and washed another three times with TBS-T. Blots were imaged using an Odyssey infrared imager.

### Yeast two-hybrid

Plasmids containing the Mob179-314 and Mob245-278 were fused to Gal binding domains and transformed into the Y2H Gold strain. Dbf2 and Cbk1NTR were fused to the Gal activation domain and transformed into Y187. Serial dilutions of cultures were grown on –Leu –Trp –His triple dropout media containing 25mM 3-Amino-1,2,4-triazole (3-AT) until colonies were visible.

### Molecular dynamics simulations

MD simulations were carried out as described earlier but using the revised Cbk1-Mob2 crystal structure presented in this study (46). Since the αC of Cbk1 was not resolved in the crystal structure, we used homology modeling based on the structure of activated PKB (PMID: 12434148) (48) to build this region. Furthermore, the activation loop segment with the Ser-570 autophosphorylation site as well as the DFG loop of the Cbk1-Mob2 complex was remodeled to make it adopt a similar structure as it had formerly been observed in other activated AGC kinases.

### Structure deposition

The crystallographic models of the Dbf2^NTR^ -Mob1, Cbk1^NTR^ -Mob2 and the revised Cbk1-Mob2-pepSsd1 complexes have been deposited in the Protein Data Bank with accession codes: 5NCN, 5NCM and 5NCL, respectively.

## Acknowledgments

We acknowledge a grant of computer time from CSCS Swiss National Supercomputing Centre, and NIIF Hungarian National Information Infrastructure Development Institute.

## Supplementary Figures

**Supplementary Figure 1.**
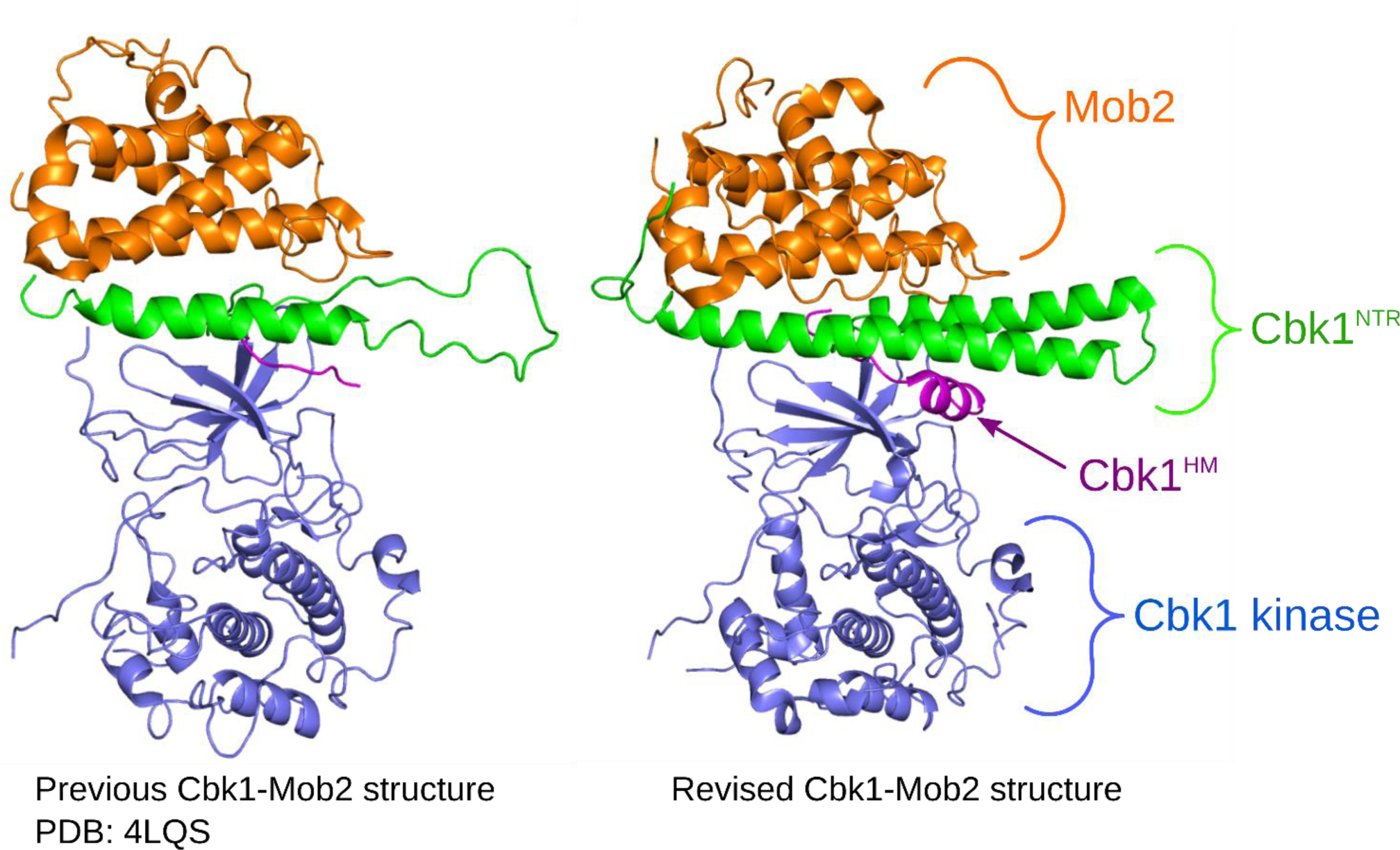
Revised full-length Cbk1-Mob2 structure. Crystallographic data from the previous Cbk1-Mob2 structure was refined using the crystallized 2.8Å zinc-binding Cbk1^NTR^-Mob2 structure. Left, the previously published full-length Cbk1-Mob2 structure (PDB: 4LQS); right, the revised crystal structure (PDB: 5NCL). In particular, the Cbk1^NTR^ (green) was re-threaded, forming a bi-helical bundle, and more closely matches other Ndr/Lats-Mob structures. Also visible is the Cbk1 hydrophobic motif (Cbk1^HM^) which forms a helix in the pocket between the NTR and kinase domain (right, magenta). Mob2 is shown in orange; the catalytic domain is shown in blue.

**Supplementary Figure 2.**
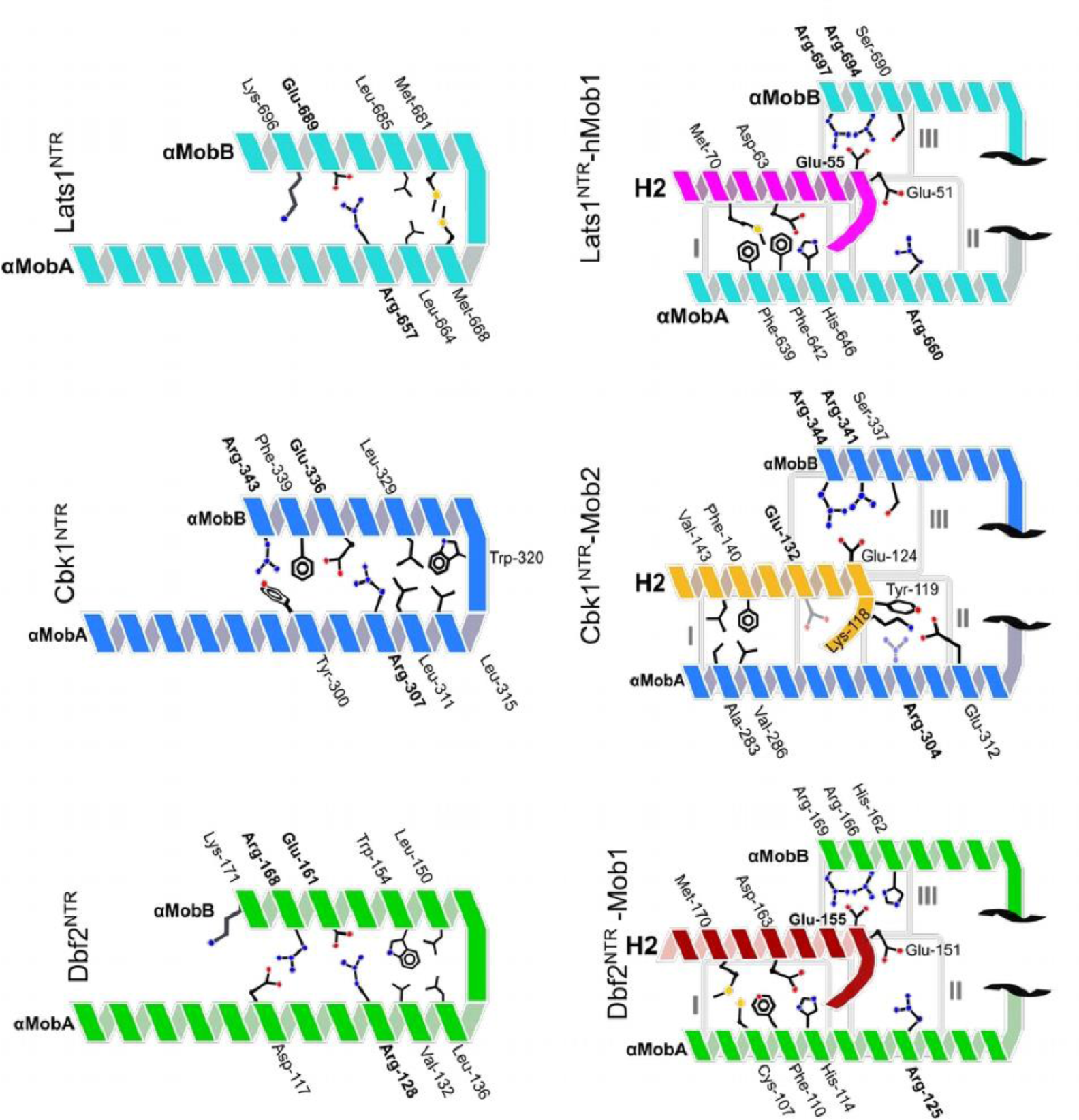
Layout of the Lats1-hMob1 interaction surface. Cartoon of primary interacting residues from Lats1^NTR^ -hMob1 (PDB ID: 5BRK) and Dbf2 ^NTR^-Mob1 compared with the revised Cbk1^NTR^ -Mob2. Conserved residues addressed in this study are bolded. Lats1, cyan; hMob1, magenta; Cbk1, blue; Mob2, orange; Dbf2, green; Mob1, dark red. Conserved interfaces I-III are boxed.

**Supplementary Figure 3.**
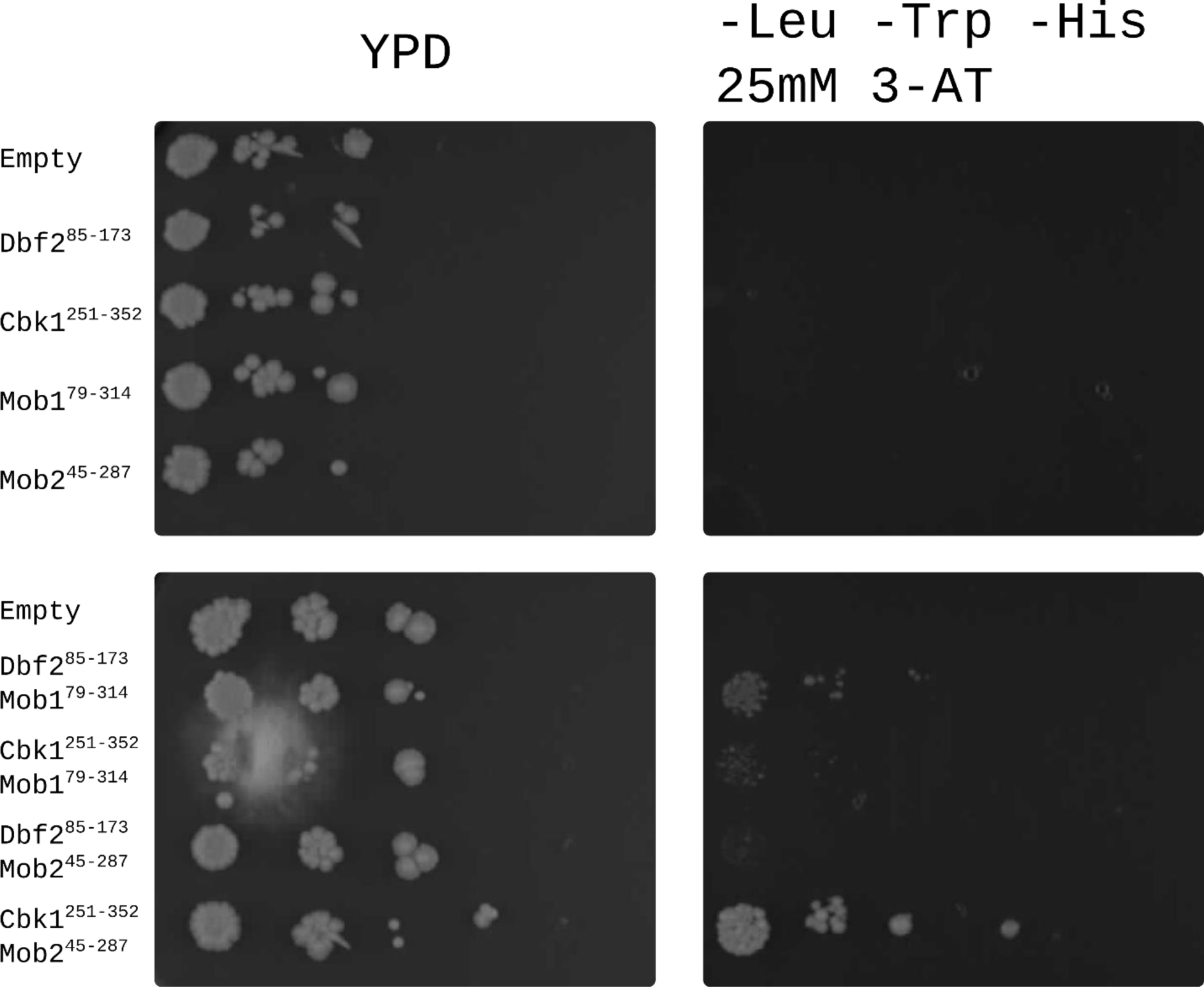
Mob-NTR specificity is maintained *in vivo.* Yeast Two-Hybrid of Kinase^NTR^/Mob fragments. Gal binding domain-Mob1^79-314^ and Mob2^45-278^ fusions were co-expressed with Dbf2/Cbk1^NTR^ fused to the Gal activation domain. Yeast cells containing protein fragments were grown on YPD or -Leu -Try -His dropout agar and serially diluted. Agar contained 25mM 3-AT to prevent autoactivation.

**Supplementary Figure 4.**
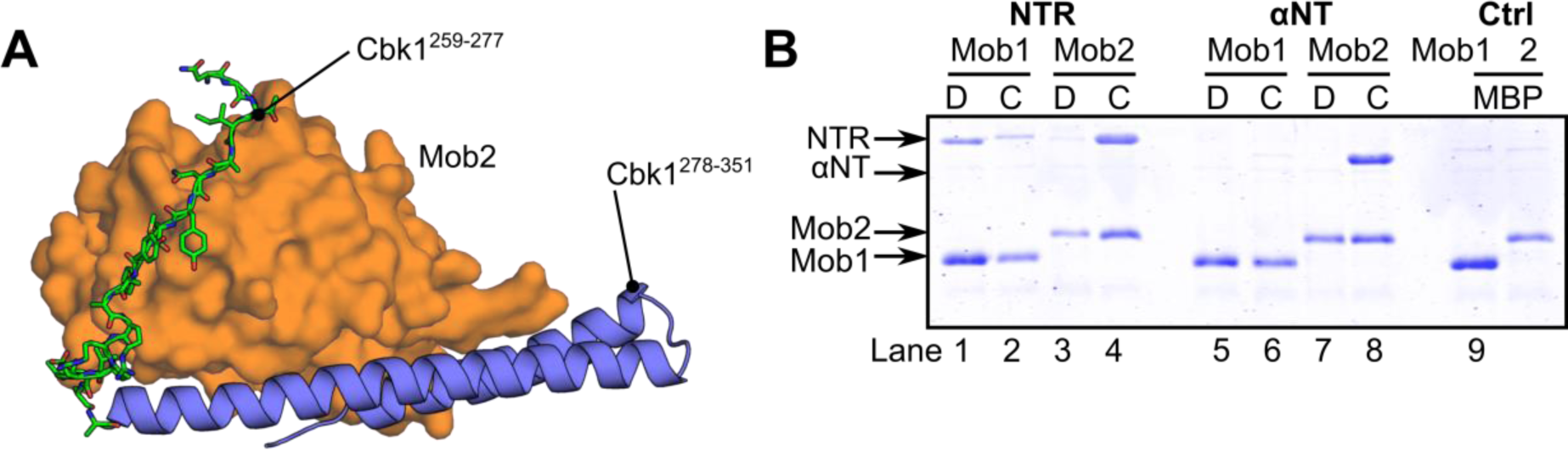
Cbk1 forms a yeast-specific binding interface with Mob2. (A) Cbk1^259-277^ (green, sticks) wraps around the back of Mob2 (orange) and potentially forms a yeast-specific Mob2-Cbk1 interface and forms the αNT along with the αMobA helix. (B) Full gel from Figure 6. Dbf2 αNT (85–124) and Cbk1 αNT (251–303) were co-expressed with Mob1 zinc-binding (lanes 5-8), and complexes isolated using IMAC as described in that figure.

**Supplementary Figure 5.**
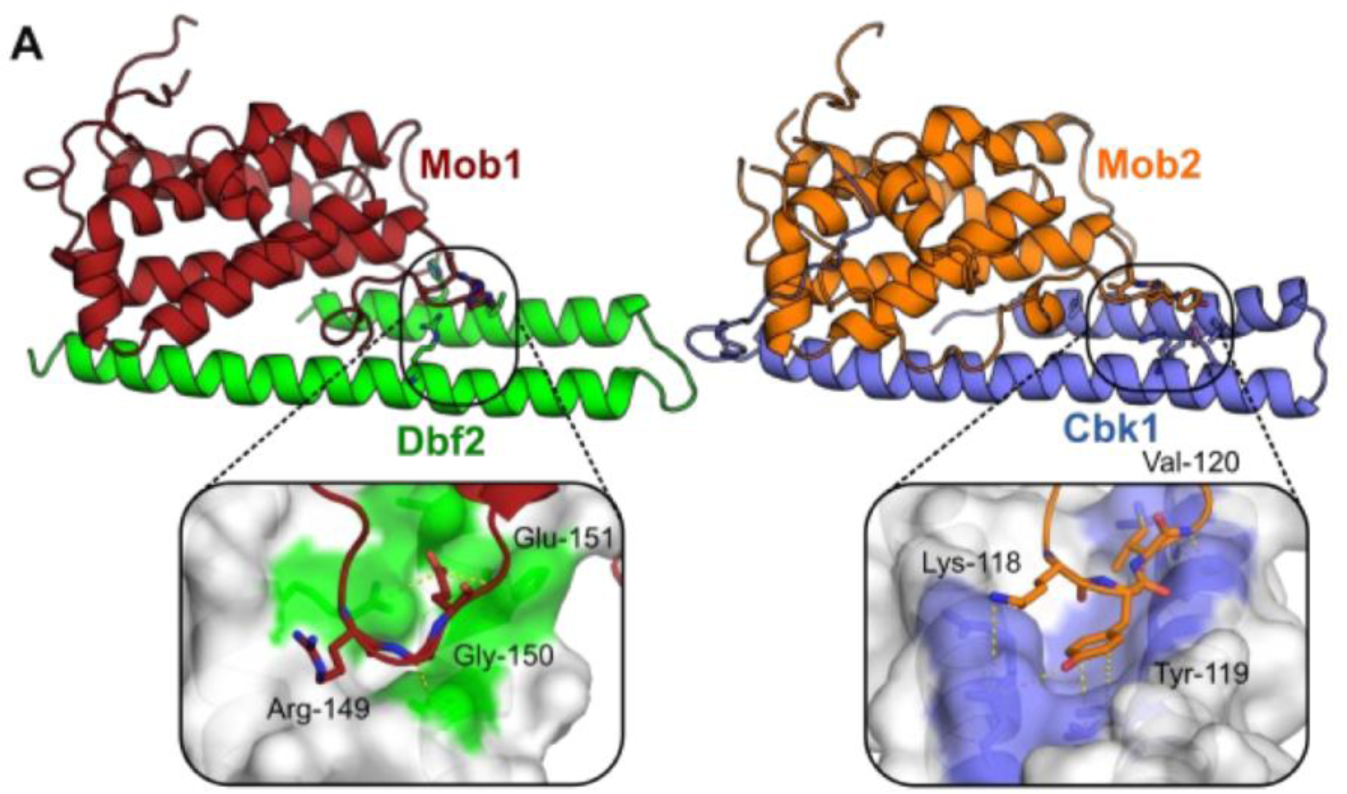
Mob1/2^KRM^ binds Dbf2 and Cbk1 through distinct binding clefts. Mob1 (dark red) binds Dbf2 (green) through the KRM tripeptide Arg-Gly-Glu. Mob2 (orange) binds Cbk1 (blue) through the KRM tripeptide Lys-Tyr-Val. Both lay in binding clefts with unique conformations formed at the junction between the NTR helices (shown as surfaces). Interactions are shown as dashed lines.

**Supplementary Figure 6.**
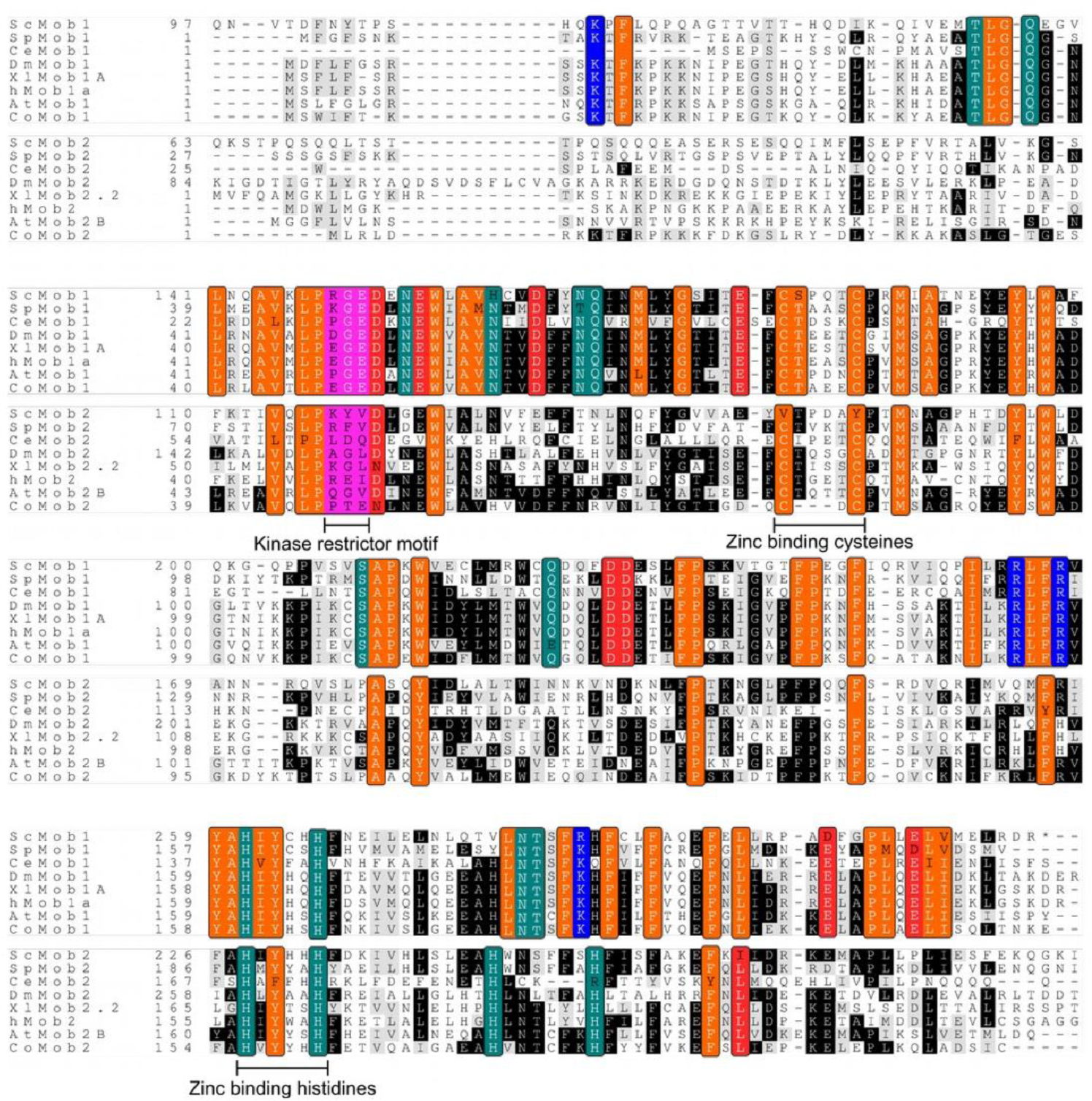
Alignments of Mob core domains. Alignment of eukaryotic Mob core domains. Sc, *Saccharomyces cerevisiae*; Sp, *Schizosaccharomyces pombe;* Ce, *Caenorhabditis elegans*; Dm, *Drosophila melanogaster*; Xl, *Xenopus laevis*; Co, *Capsaspora owczarzaki*; At, *Arabidopsis thaliana;* h, *Homo sapiens.* The KRM is highlighted in magenta; conserved hydrophobic residues, orange; basic residues, blue; acidic residues, red; polar uncharged residues, dark green.

**Supplementary Figure 7.**
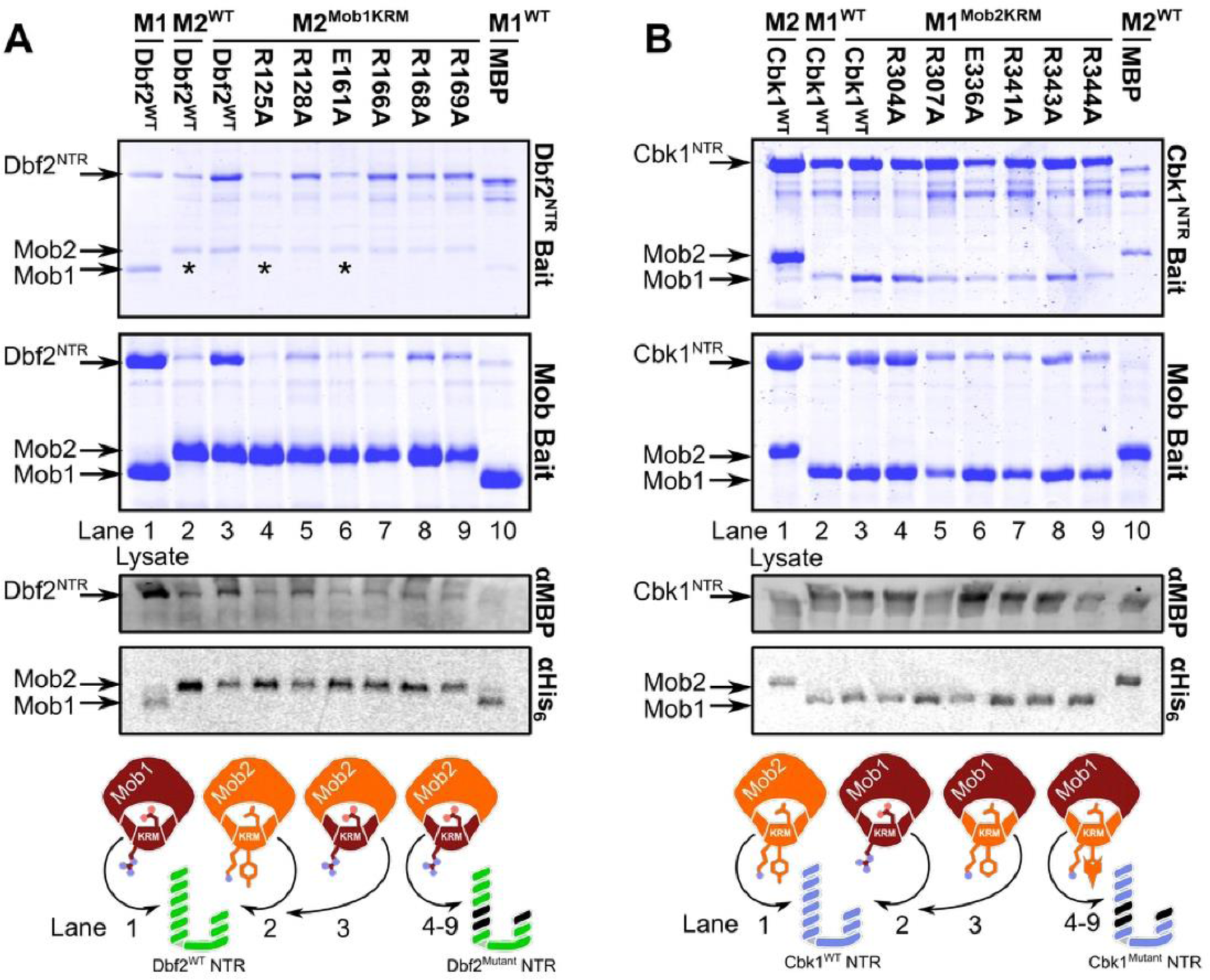
Chimeric kinase-Mob complexes bind targets through conserved residues. Chimeric kinase-Mob complexes containing swapped KRMs interact through conserved residues. (A) The Dbf2-Mob2 ^Mob1KRM^ chimera interacts through interfaces II and III. We co-expressed MBP-fused wild-type and mutant Dbf2^NTR^ with wild-type and chimeric Mob1 and Mob2 in BL21(DE3) RIL cells. We then isolated the complexes using amylose (top) and nickel chromatography (bottom), separated them using SDS-PAGE, and stained them with Coomassie. Lane 1, Dbf2^WT^-Mob1 ^WT^; 2, Dbf2 ^WT^ -Mob2 ^WT^; 3, Dbf2 ^WT^ -Mob2 ^Mob1KRM^; lanes 4-9, Dbf2^Mutant^ -Mob2^Mob1KRM^; lane 10, MBP-Mob1 ^WT^. (B) Cbk1 and Mob2/chimeric Mob1 ^Mob2KRM^ expressed and purified similarly. Lane 1, Cbk ^WT^ -Mob2 ^WT^; 2, Cbk1 ^WT^ - Mob1 ^WT^; 3, Cbk1 ^WT^ -Mob1 ^Mob2KRM^; lanes 4-9, Cbk1^Mutant^- Mob1 ^Mob2KRM^; lane 10, MBP-Mob2 ^WT^. Mob2’s strong signal in lane 10 (MBP-Mob2 ^WT^) may be due to electrostatic interactions between basic Mob2 and acidic MBP. All Mob2 constructs are stabilized by the Mob2^V148C Y153C^ mutation. Mutants: M1 ^WT^, Mob1; M2 ^WT^, Mob2; M1 ^Mob2KRM^, Mob1^R149K G150Y E151V^; M2^M1KRM^, Mob2^K118R Y119G V120E^. Constructs marked (*) adhere poorly to the resin. Brightness and contrast of the Western-blot images were adjusted as needed.

**Supplementary Figure 8.**
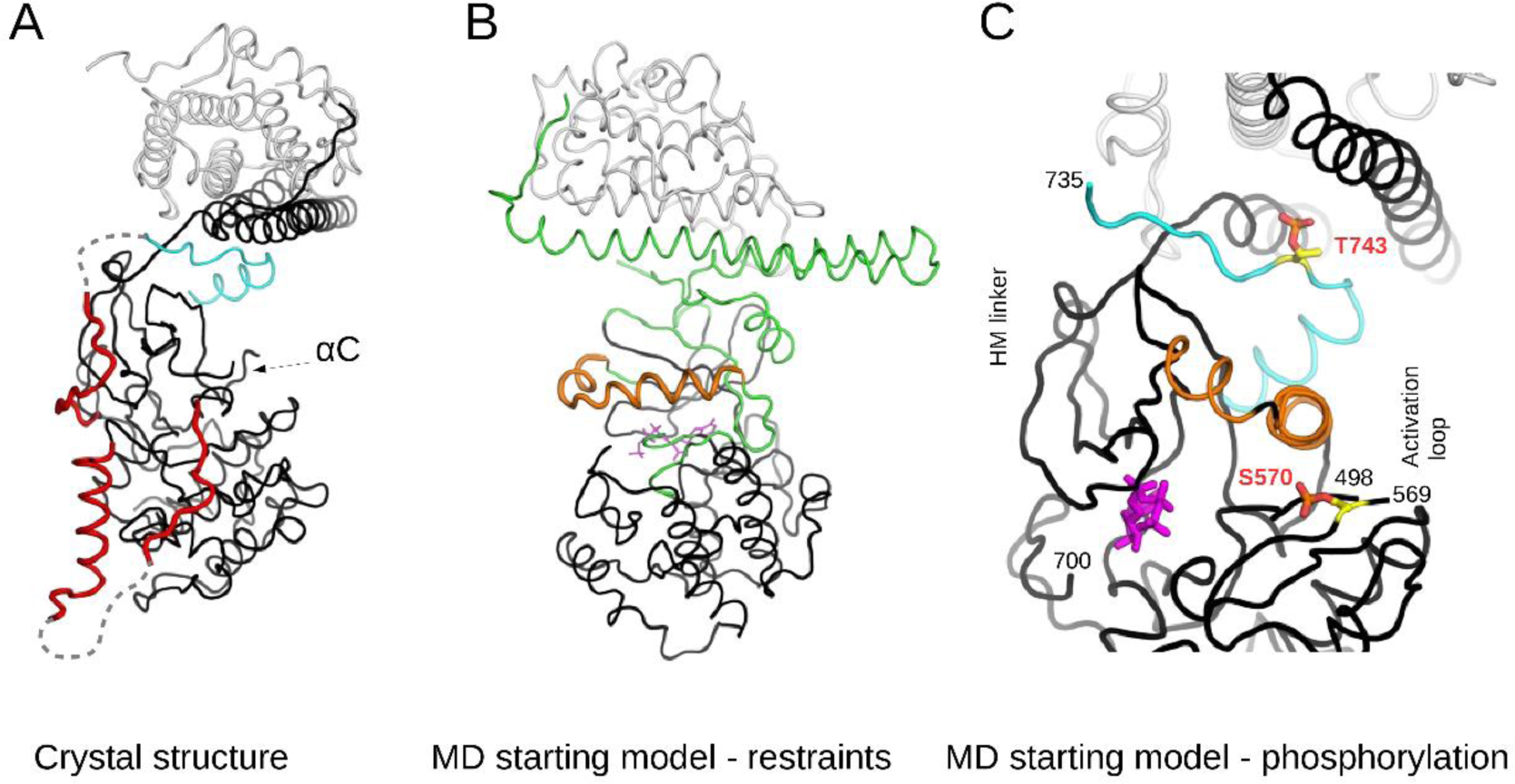
Details of MD modeling. (A) Crystal structure of the truncated Cbk1^NTR^-Mob2 complex. Regions that were not part of the MD model, because their conformation in the crystal structure was not compatible with the active state (e.g. the inhibitory helix part of the long activation loop which directly blocks the substrate binding pocket) or are parts of flexible loops (e.g. N- and C-terminal segments of the activation loop or the linker between HM and the kinase core), are colored in red. “αC” highlights that this region could not be traced in the crystal structure and is likely flexible. Dashed lines show the missing parts of flexible loops. The kinase domain core, the HM and Mob2 are colored in black, cyan and gray, respectively. (B) The starting model used for MD simulations. The hybrid model was generated using the Cbk1-Mob2 crystal structure but the conformation of DFG loop, αC and the short C-terminal activation loop segment containing the auto-phosphorylation site were all modeled based on active PKB/Akt crystal structures. Regions that were allowed to move freely are colored in green. The homology modeled αB + αC region (385–406) is colored in orange and main chain atom restraints had to be applied to keep this region in helical conformation. Further parts of the model with heavy atom restraints were the following: 1) Cbk1: 352-382, 423-468, 478-492, 569-700; 2) the full Mob protein and 3) the adenosine ring of the ATP. The ATP nucleotide is shown in magenta. (C) The model of the active Cbk1-Mob2 complex. The panel shows the starting MD model of the double-phosphorylated complex where amino acid residues (Ser-570 and Thr-743) involved in the regulation of kinase activity are highlighted. Protein chains with artificial breaks (at 498 and 569 or at 700 and 735) were N-methylated or acetylated.

